# Upregulated PD-1 Signaling is an Important Antagonist to Glomerular Health in Aged Kidneys

**DOI:** 10.1101/2021.10.26.466006

**Authors:** Jeffrey W. Pippin, Natalya Kaverina, Yuliang Wang, Diana G. Eng, Yuting Zeng, Uyen Tran, Carol J. Loretz, Anthony Chang, Christopher O’Connor, Markus Bitzer, Oliver Wessely, Stuart J. Shankland

## Abstract

Kidney aging and its contribution to disease and its underlying mechanisms are not well understood. With an aging population, kidney health becomes an important medical and socioeconomic factor. We previously showed that podocytes isolated from aged mice exhibit increased expression of Programed Cell Death Protein 1 (PD-1) surface receptor and its two ligands (PD-L1, PD-L2). *PDCD1* transcript increases with age in micro-dissected human glomeruli, which correlates with lower eGFR, and higher segmental glomerulosclerosis and vascular arterial intima to lumen ratio. *In vitro* studies in podocytes demonstrate a critical role for PD-1 signaling in cell survival and induction of a Senescence-Associated Secretory Phenotype (SASP). To prove PD-1 signaling is critical to podocyte aging, aged mice were injected with anti-PD-1 antibody (aPD-1ab). Treatment significantly improved the aging phenotype in both kidney and liver. In the glomerulus, it increased the life-span of podocytes, but not parietal epithelial, mesangial or endothelial cells. Transcriptomic and immunohistochemistry studies demonstrate that anti-PD-1 treatment improved the health-span of podocytes. It restored the expression of canonical podocyte genes, transcription factors and gene regulatory networks, increased cellular metabolism signatures and lessened SASPs. These results suggest a critical contribution for increased PD-1 signaling towards both kidney and liver aging.

## INTRODUCTION

The US population is aging, and the numbers of Americans ages 65 and older will more than double over the next 40 years (US Census Bureau), and Eurostat Predictions forecast that 28% of Europeans would be over 65 years by 2060. As the life expectancy increases, the impact of advanced age on kidney health and function is becoming an increasingly important medical and socioeconomic factor. Glomerular Filtration Rate (GFR) declines after age 40 by 0.8-1.0% per year,(1, 2) and kidneys from 70-75 year-old healthy donors have 48% fewer intact nephrons compared to 19-29 year-old donors,(3) which is consistent with an estimated annual loss of 6,000-6,500 nephrons after age 30.(3)(4)

The characteristic physiological, histological and molecular changes to the aged kidney have been well described and reviewed.(5–10) At the cellular level, the age-dependent glomerulosclerosis and the accompanying decline in GFR are paralleled by changes in number, structure and function to all four resident glomerular cell types (*i.e.* podocytes, mesangial cells, endothelial cells and parietal epithelial cells).(11–13) By now, age-dependent glomerulosclerosis is regarded as a podocyte disorder.(14) Studies in animal models demonstrated that podocyte loss causes glomerulosclerosis in direct proportion to the degree of depletion.(15–17) Similarly, upon aging podocyte numbers and density decrease in rats and mice.(11, 18–21) In humans, Hodgin and Wiggins(22) showed that the podocyte reserve dropped about 0.9% annually from >300 per 100 cm^3^ in young kidneys to <100 per 100 cm^3^ by 70-80 years of age. In fact, older age is independently associated with both absolute and relative podocyte depletion.(23)

The causes of podocyte changes in aged kidneys are incompletely understood.(24) To gain insights into candidate mechanisms at the transcriptomic level, we recently performed bulk RNA-seq comparing podocytes from 2-3-months-old (∼20 year old human) to 24-month old mice (∼70+ year old human).(25) We were struck by the significant increase in expression of Programmed cell death 1 (PD-1, synonyms PDCD1& CD279), Programed cell death 1 ligand 1 (PD-L1, synonyms CD274 & B7-H1) and Programed cell death 1 ligand 2 (PD-L2, synonyms CD273 & B7-DC) in aged podocytes compared to young podocytes.(25) The 288 amino acid, type I membrane protein PD-1 is typically expressed on immature immune cells and is bound by the two ligands of the B7 family, PD-L1 and PD-L2.(26, 27) In addition to being expressed on circulating immune cells, PD-1, PD-L1 and PD-L2 are also expressed in various cancers and many other cell types.(28) Clinical studies have shown that blocking the PD-1/PD-ligand signaling pathway with checkpoint inhibitors improves outcomes in many forms of cancer.(29, 30)

Based on the increased mRNA levels of PD-1 and its ligands in aged podocytes, we hypothesized that reducing PD-1/PD-ligand signaling in the aged kidney would improve podocyte health. To this end, we performed both *in vitro* and *in vivo* experiments to systemically address the function of PD-1 signaling on glomerular health during aging.

## RESULTS

### PD-1 and PD-1 Ligands increase in aged mice and human kidneys

The premise for this study was our recently published mRNA-seq data that showed a 17.4-fold increase in transcript levels for PD-1, a 1.4-fold increase in PD-L1 and a 2.3-fold increase in PD-L2 in podocytes from aged mice compared to young mice.(25) This was confirmed in the current study in an independent cohort of young (n=16) and aged (n=15) mice, where mRNA expression was assayed by qPCR performed on magnetic activated cell sorted (MACS) podocyte fractions. *PD-1* increased 6.7-fold (1.6±0.2 young *vs.* 10.1±1.2 aged, p<0.0001), *PD-L1* increased 2.3-fold (3.6±0.3 young *vs.* 8.4±0.8 aged, p<0.0001) and *PD-L2* increased 17.3-fold (12.8±2.9 young *vs.* 221.4±35.7 aged, p<0.0001) in aged podocytes.

Immunofluorescent staining substantiated these findings at the protein level, showing that PD-1 was barely detected in young mouse kidneys, but was markedly elevated in aged kidneys (**Figure 1A-D**). Co-immunostaining of anti-PD1 antibody with cell type-specific markers showed that in aged mice the increased PD1 protein co-localized with Nephrin and Synaptopodin-positive podocytes and also localized to parietal epithelial cells lining Bowman’s capsule (**Figure 1B,D**). This was accompanied by increased PD-L1 staining (**Figure 1E,F**). In addition, LTL-positive proximal tubular epithelial cells (**Figure 1 G,H**) and interstitial CD45-positive lymphocytes (**Figure 1 I,J**) were PD-1 positive, while alpha 8 integrin-positive mesangial cells (**Figure 1 K,L**) and CD31-positive glomerular endothelial cells (**Figure 1 M,N**) were PD-1-negative. The same staining pattern was observed in human kidneys, where PD-1 immunostaining was also increased in aged podocytes, PECs and tubular epithelial cells, but not in young human kidneys (**Figure 1O-R**).

**Figure 1.**
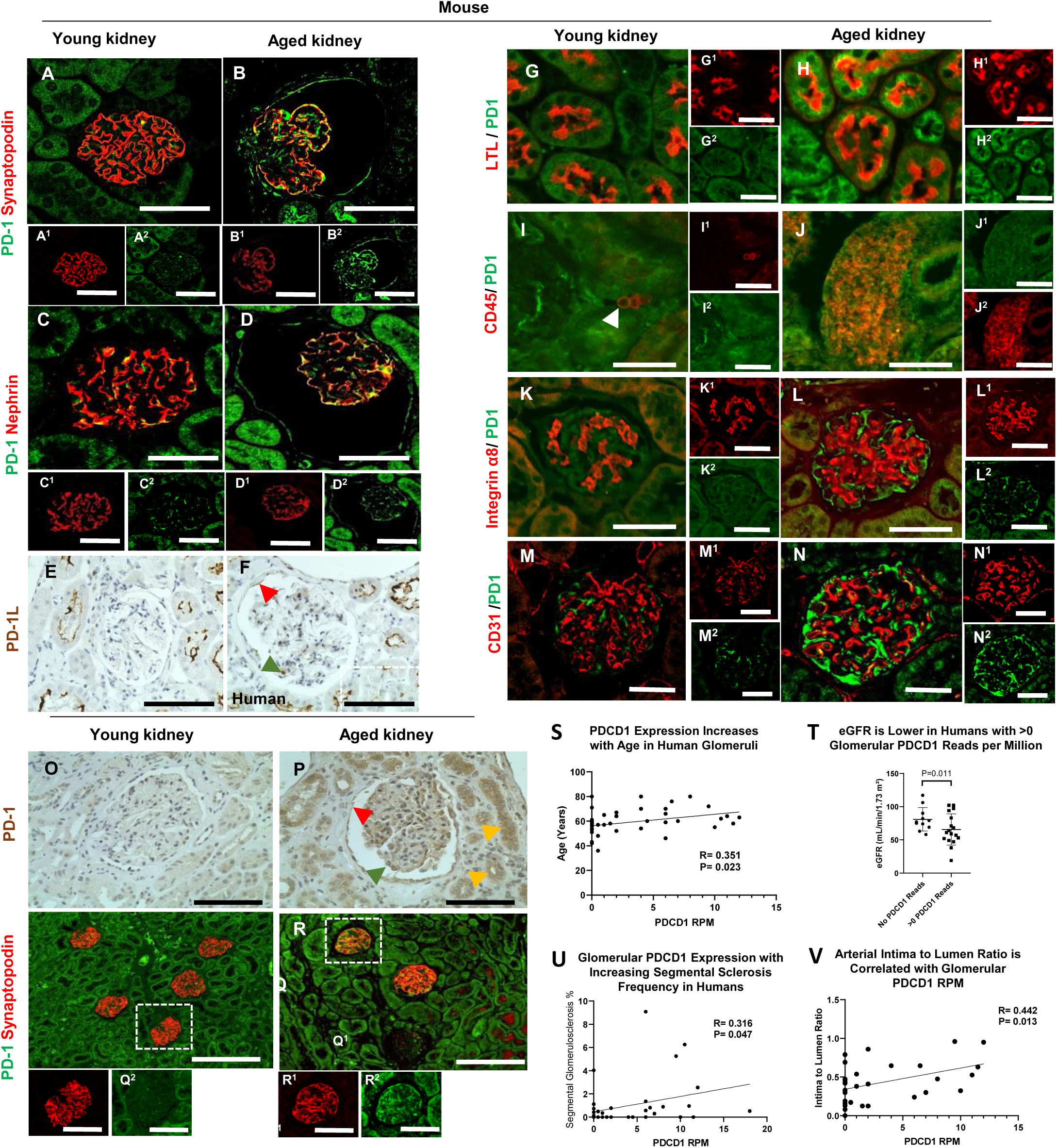
*Podocyte PD-1 Immunostaining and transcripts*. (A-N) *Mouse kidney*. Representative images of immunofluorescent stains show the merge of two antibodies. Images labeled with superscripted numbers show individual red and green fluorescent channels. (A-B) Double immunofluorescent staining of PD-1 (green) and synaptopodin (red) shows that PD-1 is not expressed in the normal mouse glomerulus (A) but merges with synaptopodin positive podocytes in the aged kidney (B, yellow color). PD-1 also stains PEC along Bowman’s capsule in the aged kidney (B, green color). (C-D) Double immunofluorescent staining of PD-1 (green) and nephrin (red) shows that PD-1 staining is absent in young glomeruli (C) and merges with the podocyte protein nephrin in the aged kidney (D, yellow). PD-1 staining is increased in PEC and the proximal tubular epithelial cells in the aged kidney (D, green). (E-F) Immunoperoxidase staining for PD-1L (brown) is not detected in young mouse kidney (E), but is detected in a podocyte (green arrow), PEC (red arrow) and proximal tubules in the aged mouse. (G,H) LTL (red) stains the brush border of proximal epithelial cells and merges with PD1 (green) in the aged kidney (H). (I,J) CD45-positive interstitial lymphocytes (red) merge with PD1 (green) in the aged kidney (J). Increased PD-1 staining (green) does not merge with the mesangial cell marker alpha 8 integrin (red) (L). Increased PD-1 (green) does not merge with the endothelial cell marker CD31 (red) in the aged kidney (N). (O-R) *Human kidney*. (O,P) Immunoperoxidase staining for PD-1 (brown) is not detected in young human kidney (O), but is present in podocytes (green arrow), PECs (red arrow) and tubular epithelial cells (orange arrows) in aged human kidney (P). (Q) Double immunofluorescent staining for PD-1 (green) and synaptopodin (red) shows no PD-1 in the young human glomerulus. (R) PD-1 staining merges with synaptopodin (yellow color) in the aged human glomerulus. (S-V) *PDCD1 transcripts from micro-dissected human glomeruli*. Expression of *PDCD1* (∼human PD-1) increased with age (S), and was accompanied by lower eGFR (T), higher segmental glomerulosclerosis (U) and higher vascular injury (V). These results show that PD-1 is increased in podocytes, PECs and proximal tubules in both mouse and human aged kidneys, and high glomerular expression in humans correlates with reduced kidney function and higher glomerular scarring.

Moreover, examining a potential functional consequence of PD-1 expression in aged humans we analyzed transcriptomic data from micro-dissected glomeruli from aged human kidneys for correlations between *PDCD*1 (∼human PD1) and clinical parameters for glomerular aging and function. *PDCD1* expression increases with human age (P<0.023, R=0.351, **Figure 1S**). Importantly the increased *PDCD1* transcript levels were accompanied by a lower eGFR (P=0.011, **Figure 1T**), and correlated with increased segmental glomerulosclerosis (P=0.047, R=0.316, **Figure 1U**) as well as reduced arterial intima to lumen ratio (P=0.013, R=0.0442, **Figure 1V**), a measure of vascular injury.

These results show that the PD-1 pathway increases in aged mouse and human glomeruli, and is a strong predictor of declining kidney function, glomerular scarring and vascular damage.

### Overexpression of PD-1 is sufficient to induce death in cultured podocytes

To address whether increased PD-1 signaling has a biological role we utilized immortalized mouse podocytes,(31–33) which endogenously expresses one of the PD-1 ligands, PD-L1, yet had very low levels of PD-1 itself and its other ligand, PD1-L2 (**Figure 2A,D,F)**. Ectopic expression resulted in highly elevated PD-1 levels but did not impact the expression of either ligand (**Figure 2A,E,G)**. Overexpression of PD-1 had dramatic effects on podocyte survival, inducing apoptosis as assessed by cleaved Caspase-3 staining and dead cell quantification (**Figure 2H-L)**. To verify that this was indeed due to induced PD-1 signaling, we used a neutralizing anti-PD-1 antibody (referred to as aPD1ab). Treating PD-1 overexpressing podocytes with aPD1ab restored the levels of podocyte death to the levels observed for the GFP vector control podocytes (**Figure 2M-Q**). Thus, *in vitro* experiments in immortalized podocytes support a critical role for PD-1 signaling in podocyte survival that could contribute to podocyte aging *in vivo*.

**Figure 2.**
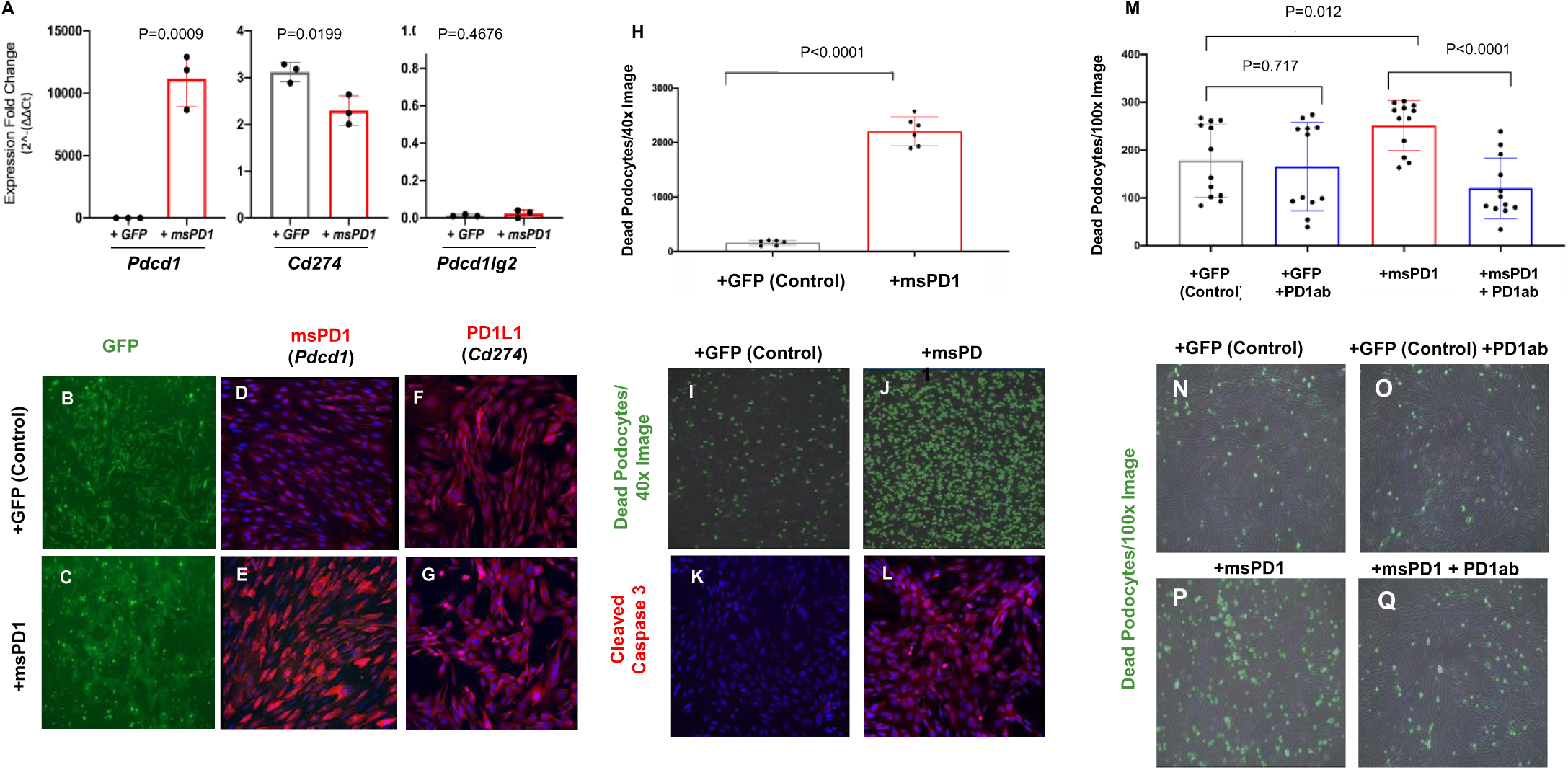
*Overexpressing PD1 induces podocyte death in cell culture*. (A) Following the overexpression of PD1 utilizing a lentiviral expression vector (msPD1, red bars) in immortalized mouse podocytes, mRNA levels increased significantly for *Pdcd1* (PD1) compared to control GFP infected podocytes, without changes to *Cd274* or *Pdcd1lg2*. GFP expression (green) of the GFP control (B) and PD1 overexpressing (C) lentiviral vectors confirm efficient transfection of immortalized mouse podocytes. Immunocytochemistry for PD1 protein (red) in GFP control infected podocytes (D), was significantly increased in msPD1 overexpressing podocytes (E). DAPI stains nuclei blue. Staining for PD1 ligand 1 (red) was not different between GFP control infected (F) and PD1 overexpressing (G) podocytes. (H) The overexpression of PD1 (red bar) increased podocyte death compared to control GFP infected podocytes (gray bar). (I,J) Representative images of dead podocytes encircled with green annotations. Cleaved caspase 3 staining (red) was barely detected in GFP control infected podocytes (K), but was markedly increased in PD1 overexpressing podocytes (L). DAPI stains nuclei blue. (M) Applying anti-PD1ab (blue bar, 2^nd^ column) did not impact cell death in GFP control infected podocytes (gray bar). The increased podocyte death induced by overexpressing PD1 (red bar) was reduced when anti-PD1 antibody was applied (blue bar, 4^th^ column). (N-Q) Representative images of dead podocytes encircled with green annotations are shown for panel M graph.

### Impact of PD-1 Inhibition on Glomerular Aging in Mice

To test whether the upregulation of PD-1/PD-L1/PD-L2 is biologically relevant for aged podocytes *in vivo*, we inhibited PD-1 signaling using a neutralizing anti-PD-1 antibody (<u>abbreviated as aPD1ab</u>). Twenty-one-month-old mice were randomized and injected intraperitoneally with aPD1ab or the control IgG2a antibody (referred to as <u>IgG2a</u>) once weekly for a total of 8 weeks (**Supplementary Figure S1A**). Uninjected 4-month-old mice were used as young age controls. Treatments had no major impact on mortality, body weight or kidney function as measured by blood urea nitrogen (BUN) and urinary albumin to creatinine ratios (ACR)(**Supplementary Figure 1B-F**). Organs (blood, kidney, liver and spleen) were extracted and processed for immunofluorescence/histology. To specifically assess the effect of aPD1ab on the podocyte transcriptome, kidneys were digested and separated into podocyte and non-podocyte cell fractions by magnetic activated cell sorting (MACS).

We next addressed whether the aPD1ab treatment impacted kidney aging. As expected, staining for senescence-associated beta-galactosidase (SA-ßgal)(8) was higher in both glomeruli and tubular epithelial cells in aged mice (**Figure 3B**) when compared to their young counterparts (**Figure 3A**). Treatment with aPD1ab caused a dramatic reduction in both glomerular and tubular SA-ßgal (**Figure 3C**). The same pattern was observed in immunostainings for the senescent proteins p16 and p19,(8) where aPD1ab-injected mice exhibited reduced staining compared to age-matched control kidneys (**Figure 3D-I**).

**Figure 3.**
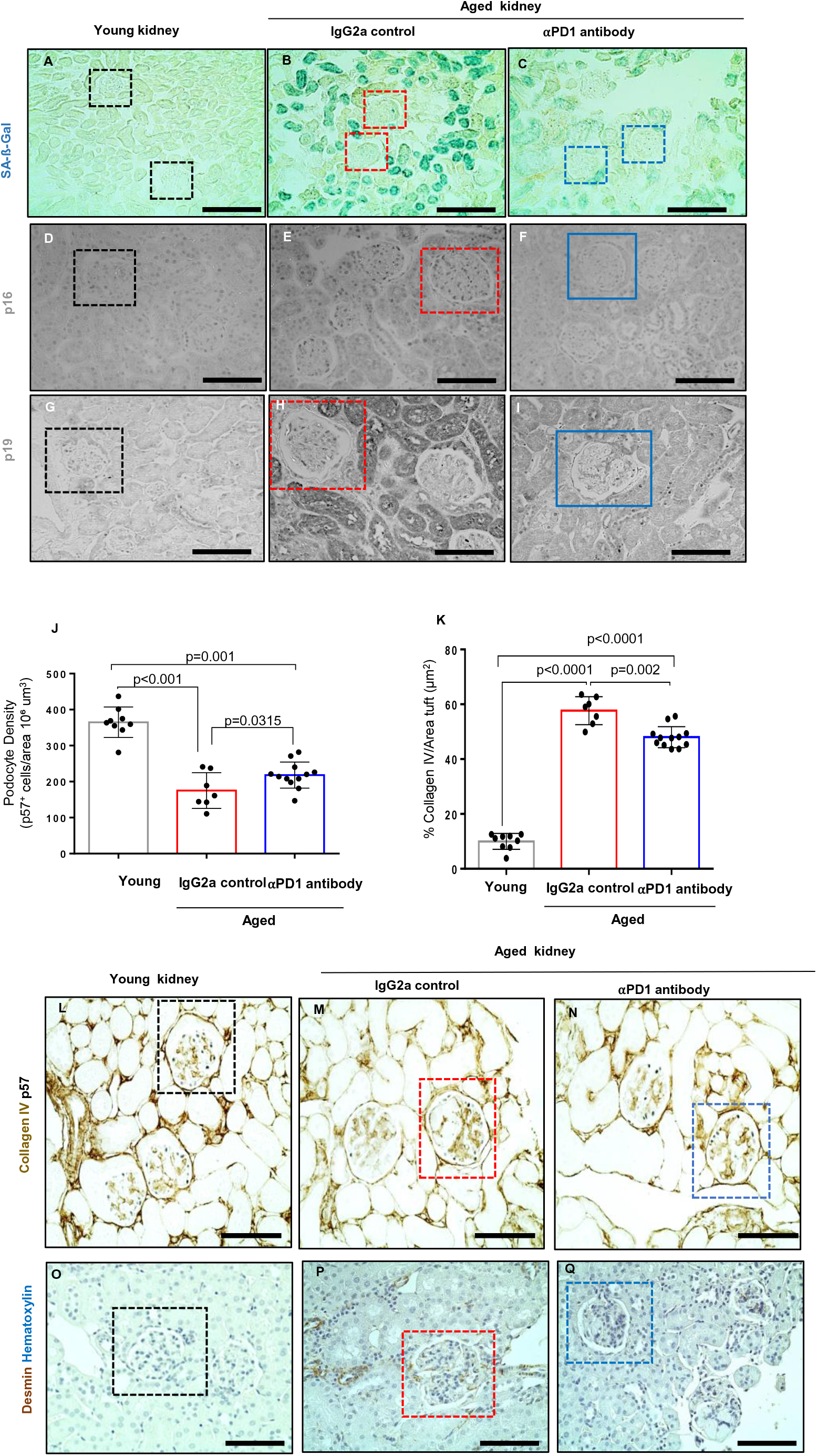
*Podocyte senescence, density and stress*. (A-I) *Senescence markers*. (A-C) Representative images of SA-ß-Gal staining (blue). SA-ß-Gal was barely detected in young kidneys (A), but increased in glomeruli (red boxes) and tubular epithelial cells in IgG2a injected aged mice (B). aPD1ab lowered SA-ß-Gal in glomeruli (blue boxes) and tubules in aged mice (C). (D-F) p16 staining (black). p16 is occasionally detected in glomeruli and tubular epithelial cells of young kidneys (D). Staining for p16 increased in glomeruli (red box) and tubular epithelial cells in the glomeruli and tubular epithelial cells of IgG2a injected mice (E) but was lower in aPD1ab injected mice (F). (G-I) p19 staining (black). p19 was barely detected in young kidneys (G) but was increased in glomeruli (red box) and tubular epithelial cells in IgG2a injected mice (H). p19 is lower in aPD1ab injected mice (I). Podocyte density was lower in aged IgG2a injected mice compared to young mice, and was increased in aged aPD1ab injected mice. (J) Each circle represents an individual mouse. Glomerular collagen IV staining was higher in IgG2a injected aged mice compared to young mice. aPD1ab lowered glomerular collagen IV immunostaining. Each circle represents an individual mouse (K). (L-M) Representative images of immunoperoxidase staining for the podocyte marker p57 (blue) and collagen IV (brown). (O-Q) Immunoperoxidase staining (brown) for the podocyte stress marker desmin was increased in aged IgG2a injected mice (P), and was lower in aged aPD1ab mice (Q). These results show that podocyte density is higher and podocyte stress lower in aged mice given aPD1ab.

A second hallmark of glomerular aging is the decrease in the podocyte’s life-span, measured by a decrease in podocyte density.(22) Measuring podocyte density using a computer-assisted machine learning approach showed a decrease in aged (*i.e.*, IgG2a-control injected) *vs.* young mice (p<0.001), which was partially restored in the aged-matched mice injected with aPD1ab (p=0.0315) (**Figure 3J**). This was accompanied by changes in glomerular collagen IV(21) (**Figure 3L-M**) and the stress marker Desmin(14, 18) (**Figure 3O-Q**). Both were increased in aged mice and reduced upon treatment with the aPD1ab.

We next analyzed the other resident glomerular cell types (**Figure 4**). Parietal epithelial cells (PECs) marked by the expression of Pax8 were decreased with age, but in contrast to podocytes, their numbers were not restored by aPD1ab treatment (**Figures 4A-D**). However, another aspect of PEC aging, the induction of an activated PEC phenotype as measured by increased expression of the activation markers CD44, CD74, pERK and an increase in Collagen IV staining along Bowman’s capsule(20, 21) were significantly restored in PECs of aged mice injected with aPD1ab (**Figures 4E-P**).

**Figure 4.**
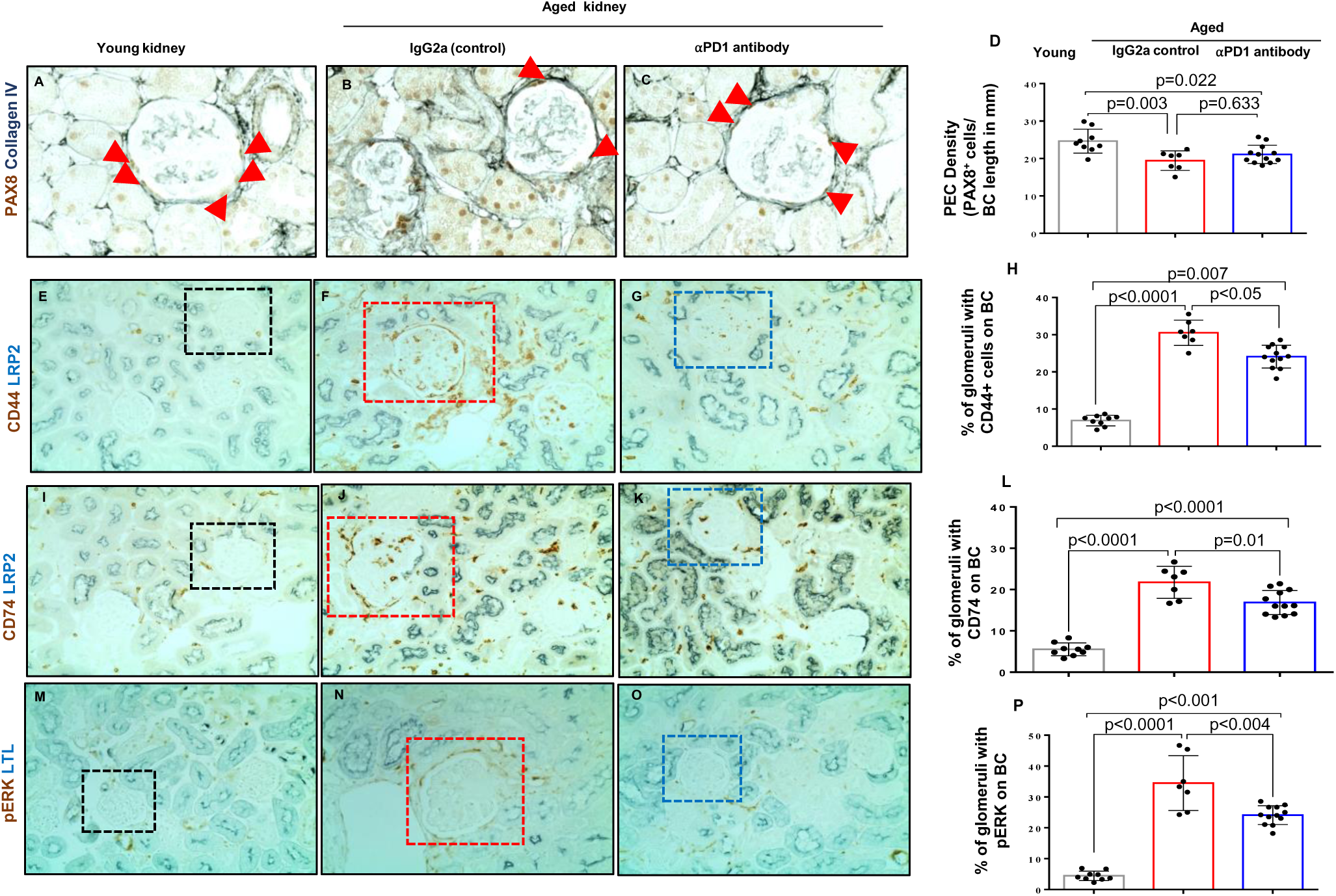
*Parietal Epithelial Cell (PEC) changes in aPD1ab injected mice*. (A-C) Representative images of immunoperoxidase staining for the PEC marker PAX8 (brown) and collagen IV (blue, outlines Bowman’s capsule). (D) PEC density was lower in aged IgG2a injected mice (red bar) compared to young mice (gray bar) but did not change with aPD1ab treatment (blue bar). Representative images of immunoperoxidase double staining with antibodies to the PEC activation markers (brown) CD44 (E-G), CD74 (I-K) and pERK (M-O) with the proximal tubular cell markers LRP2 (blue) and LTL (blue). (H,L,P) All 3 PEC activation markers were higher in aged IgG2a injected mice (red bars) compared to young mice (gray bars). aPD1ab lowered all 3 PEC activation markers in aged mice (blue bars).

The same scenario was observed for glomerular endothelial cells (GEN). GEN density, identified by nuclear staining for the ETS Transcription Factor ERG (ERG)(34, 35) was reduced in aged IgG2a control mice (443±10 young *vs.* 290±19 aged IgG2a control ERG^+^ nuclei x 10^6^ um^3^, p<0.0001), but was unchanged with aPD1ab (290±19 aged IgG2a control *vs* 287±10 aged aPD1ab ERG^+^ nuclei x 10^6^ um^3^, p=0.902) (**Supplementary Figure 2A-D**). Yet, their age-dependent increase in the fenestral diaphragm protein plasmalemmal vesicle associated protein-1 (PV-1), normally absent in healthy GEN and when present, represents an immature and injured phenotype,(36–38) was lowered by administration of aPD1ab (**Supplementary Figure 2E-G**). Finally, mesangial cell area stained by Itga8 (Integrin alpha-8) was increased in aged IgG2a control mice compared to young mice, a trend, that was also not changed by aPD1ab (**Supplementary Figure 2H-K**).

Together, these data demonstrate that interfering with PD-1 signaling can partially reverse the glomerular aging phenotype and improve podocyte life-span both in respect to cell number and function. While it did not impact the numbers/life-span of the other glomerular cell types, it did have beneficial impacts on age-dependent activation of PEC and GEN.

### Transcriptomic Changes in Aged Podocytes Modified by anti-PD1 antibody

To understand the underlying molecular mechanism of PD-1 inhibition, we wondered whether this was due to changes in the mRNA levels of PD-1 and its ligands. In podocytes, aPD1ab did not alter the mRNA expression of PD-1 and PD-L1 measured by qRT-PCR, but lowered PD-L2 (**Supplementary Figure 3 A-C**). In the non-podocyte fraction, aPD1ab lowered PD-1, but did not change levels of PD-L1 and PD-L2. mRNA levels of other members of the B7 ligand family (CD80/B7-1 and CD86/B7-2) did not change either in aged mouse podocytes (not shown). Finally, consistent with at least partial suppression of the PD-1 pathway *in vivo* several proteins in the PD-1 signaling pathway were reduced by aPD1ab (**Supplementary Figure 3 D**).

Based on this data and to obtain a better understanding of the mechanism(s) responsible for the reversal of glomerular aging by aPD1ab treatment, we performed mRNA-seq analyses from podocyte and non-podocyte cell fractions from each individual mouse from each of the groups (young, aged IgG2a-injected control and old aPD1ab-injected). Principle component analysis showed excellent clustering of the individual treatment groups (**Supplementary Figure 4 A&B**). A volcano plot identified a large number of differentially expressed transcripts in aged aPD1ab-injected *vs.* aged IgG2a control-injected podocytes (**Figure 5A**). A total of 1,137 genes were down-regulated and 949 were up-regulated in aged aPD1ab-injected compared to aged IgG2a-injected podocytes (**Figure 5B**). We next analyzed significantly altered transcripts for their contribution to the aging process. Of the down-regulated transcripts, about half (i.e. 553) genes were also up-regulated in aged *vs.* young podocytes (p=0). Similarly, of the up-regulated transcripts about a third (i.e. 300) were down-regulated in aged *vs.* young podocytes (p=1x10^-32^). 649 genes were up-regulated in aged aPD1ab injected podocytes compared to aged IgG2 injected podocytes. Together, these data suggest that more than 40% of the genes regulated by PD-1 are part of the natural podocyte aging process.

**Figure 5.**
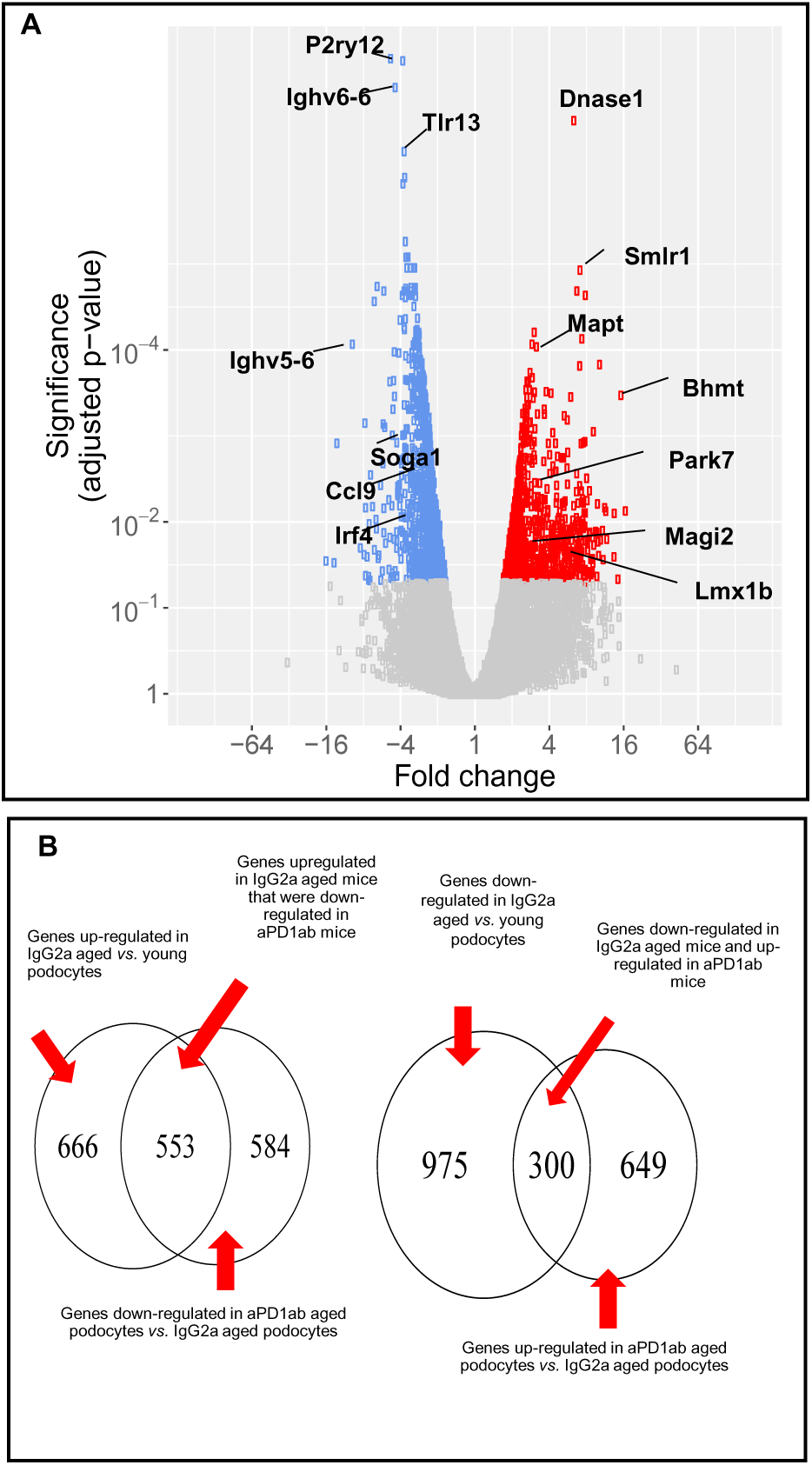
*Podocyte transcripts in aged mouse podocytes altered by anti-PD1 antibody (aPD1ab)*. (A) Volcano plot shows transcripts in aged podocytes that were decreased (blue circles), increased (red circles) or not changed (grey circles) by treating mice with aPD1antibody for 8 weeks. (B) Summary of the number of genes upregulated and downregulated in podocytes from aged control IgG2a mice compared to young mice, and from aged aPD1ab injected mice compared to age matched IgG2a injected mice.

### Podocyte Genes, Function and Transcription Factors are Restored upon anti-PD-1 Antibody-Treatment

The health-span of a podocyte can be assessed by changes to their molecular, cellular and transcriptional landscape required for their normal physiology, structure and function. To better understand how PD-1 signaling contributes to podocyte aging and how PD-1 might impact podocyte health-span, we initially focused on canonical genes essential for the highly specialized structure and function of podocytes. We have previously reported that several of these genes were decreased with aging.(39) This was confirmed with the current study cohort. More importantly, the expression of many canonical podocyte genes (e.g., *Actn4, Cdkn1, Col4a, Fat1, Lamb2, Nphs1, Nphs2, Podxl* and *Synpo*) were reduced in aged IgG2a injected mice, but significantly up-regulated upon injection with the aPD1ab (**Figure 6A**). This was confirmed by immunostaining for podocyte structural proteins nephrin (*Nphs1)* and synaptopodin (*Synpo)* (**Figure 6B-I**).

**Figure 6.**
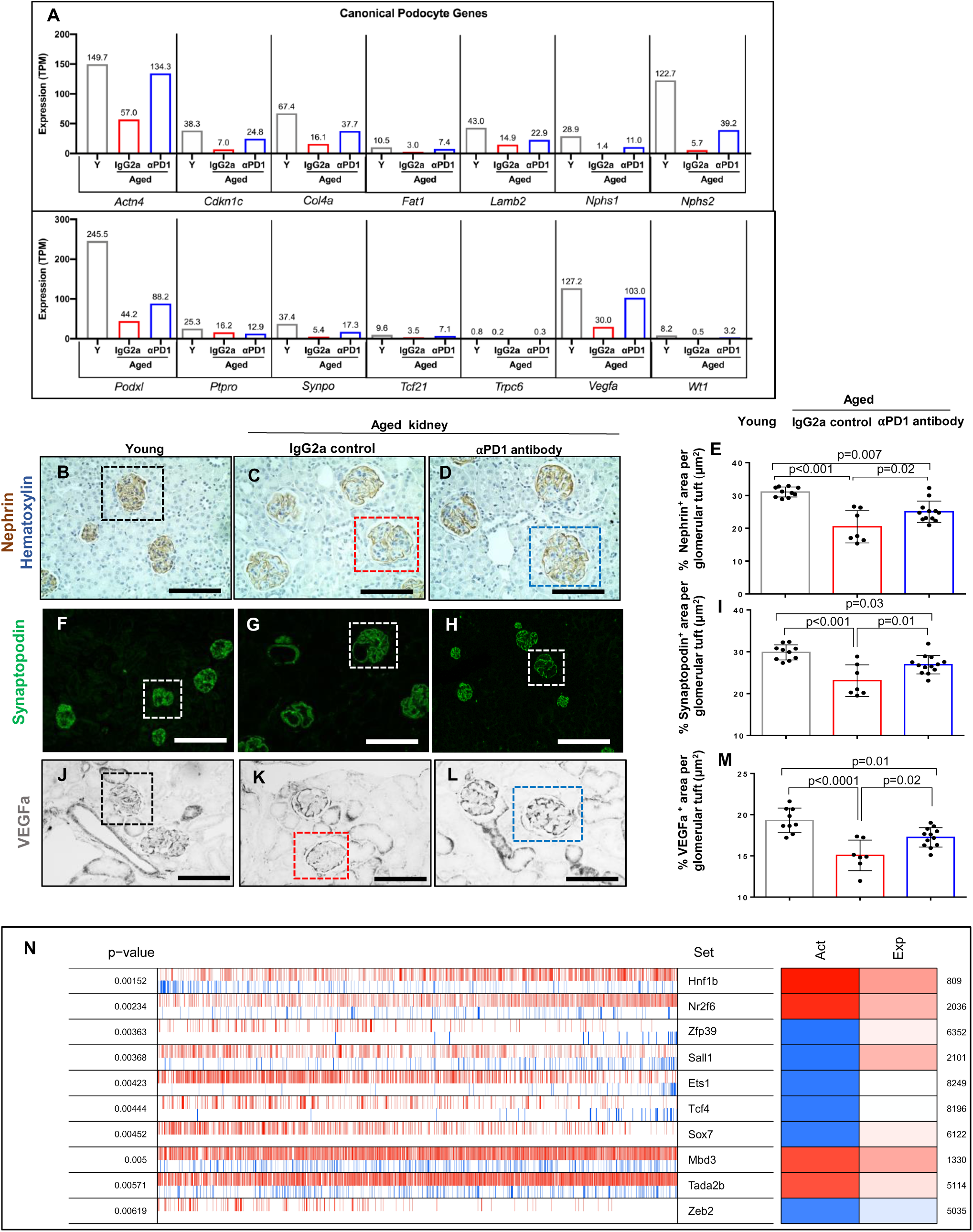
*Changes to podocyte canonical genes, proteins and transcription factors*. (A) Expression of individual Canonical Podocyte Genes in alphabetical order from RNAseq data in young mice (Y, gray bars), aged mice injected with IgG2a (red bars) and aged mice injected with anti-PD1 antibody (aPD1, blue bars). The levels of all genes that were lower in aged IgG2a injected mice compared to young mice were higher in aPD1ab injected mice, except *Ptpro*. (B-M) Protein validation of select genes by immunostaining. Representative immunoperoxidase staining for nephrin (brown) (B-D), immunofluorescent staining for synaptopodin (green)(F-H) and immunoperoxidase staining for VEGFa (black)(J-L). The boxes in each panel show an example of a glomerulus. (E, I, M) Quantitation of nephrin (E), synaptopodin (I) and VEGFa (M) staining. Compared to young mice (grayt bars), immunostaining was lower for each in aged IgG2a injected mice (red bars), but was higher in the aged aPD1ab injected mice (blue bars). Each circle represents an individual mouse. (N) VIPER (<u>v</u>irtual inference of <u>p</u>rotein activity by <u>e</u>nriched regulon) analysis of transcription factor activity. The activity of the top 10 transcription factors that were impacted by aPD1ab treatment are shown in third column, their significance (first column), representative activity (second column), conferred activity (third column) and expression (fourth column). The conferred activity of *HNf1b*, *Nr2f6*, *Mbd3* and *Tada2b* increased, with increased expression. *Zfp39*, *Sall1* and *Sox7* activity were decreased despite higher expression levels. The lower activity of *Ets1* and *Tcf4* were independent of expression levels. The activity and expression of *Zeb2* were lowered in the aPD1ab injected mice.

We also analyzed VEGFa, as a surrogate for podocyte synthetic function, which is critical for maintenance of the podocyte/endothelial cell interaction.(40) *Vegfa* mRNA levels were 4.2-fold lower in aged glomeruli than in young glomeruli, but was 3.4-fold higher in aged mice, aPD1ab-injected glomeruli (**Figure 6A**). The decreased *Vegfa* mRNA expression was validated at the protein level by immunostaining (**Figure 6J-M**). The same was observed for another pathway critical for podocyte function, tight junction formation. Again, many of the genes involved in tight junction formation were restored in the podocytes of aged mice injected with the aPD1ab compared to the IgG2a-injected controls (**Supplementary Figure 5**).

Finally, we investigated the transcriptional regulation of podocytes. Compared to control aged IgG2a-injected mice, mRNA-seq demonstrated that several podocyte transcription factors were expressed at significantly higher levels in aged mice injected with aPD1ab. These included *Lmx1b* (4.3-fold), *Osr2* (7-fold) and *Foxc2* (2-fold). The restoration of the podocyte gene regulatory network was further underscored by a VIPER (<u>v</u>irtual inference of <u>p</u>rotein activity by <u>e</u>nriched regulon) analysis. This identifies the transcription factors activities based on the expression of their downstream targets.(25, 41) As shown in **Figure 6N** from the top ten active transcription factors identified by VIPER (sorted by permutation p-value), the transcriptional activity of 4 (*i.e. Hnf1b, Nr2f6, Mbd3* and *Tada2b*) were increased, while 6 were decreased (*i.e. Zfp39, Sall1, Ets1, Tcf4, Sox7, Zeb2*) upon aPD1ab injection.

Taken together this data shows that the expression levels of canonical podocyte genes and the function and transcriptional regulation required for their health-span are partially restored with the aPD1ab.

### PD-1 Inhibition Restores Apoptosis, Pyroptosis, Endoplasmic Reticulum Stress and Autophagy

The ectopic expression of PD-1 in podocytes *in vitro* induced high levels of apoptosis (**Figure 2H-Q**). Moreover, reduced podocyte number in aging (and disease) are due to increased cell death in the face of an inability to self-renew.(42) Indeed, activated Caspase-3 staining in podocytes was increased in aged IgG2a-injected control mice (2.59 ±1.19 P<0.0001 vs control) and reduced upon aPD1ab injection (0.66±0.87 P=0.0001)( **Figure 7A-D**).

**Figure 7.**
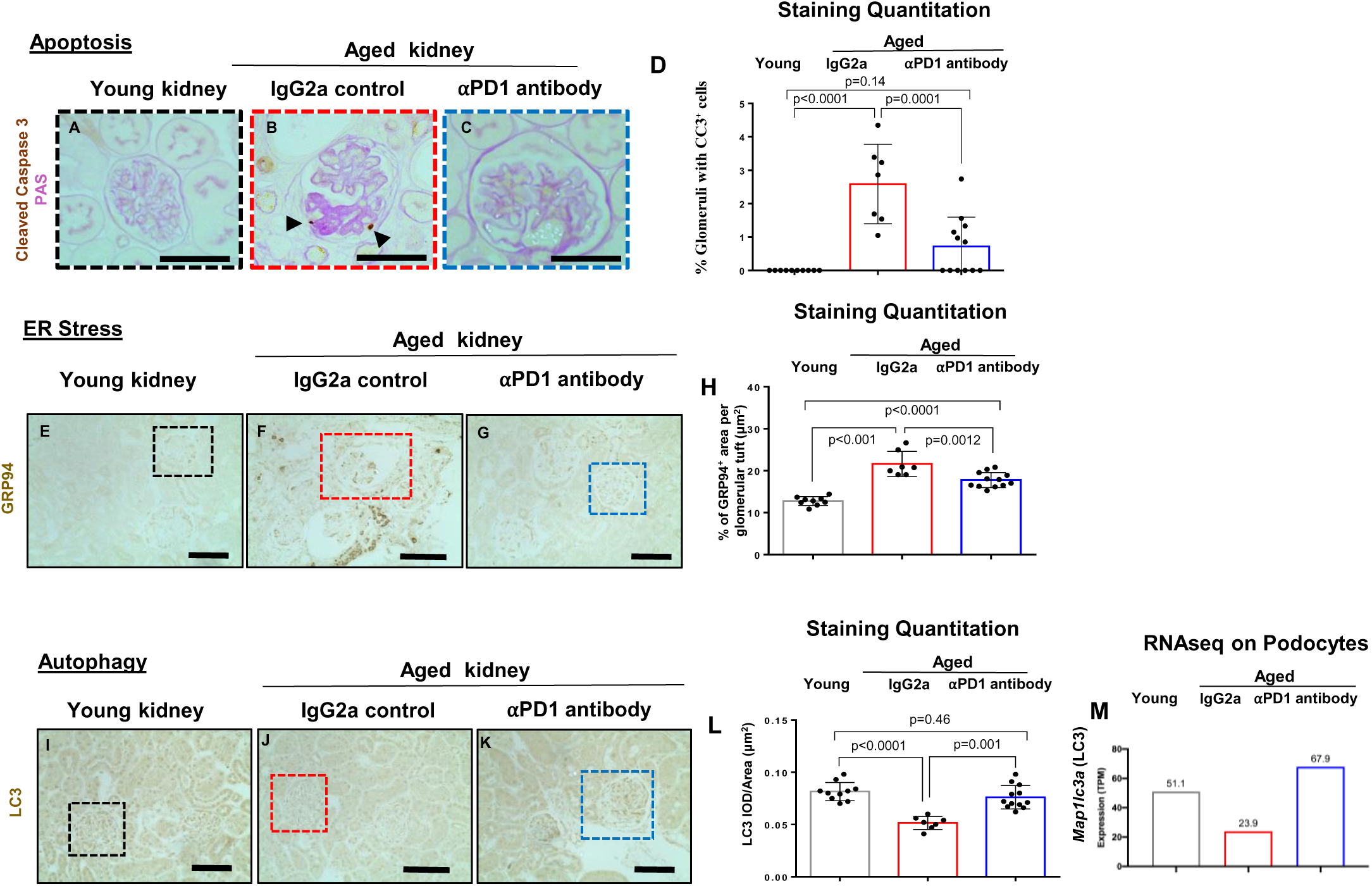
*Apoptosis, ER stress and Autophagy*. (A-D) Representative images of immunoperoxidase staining for the apoptosis marker cleaved caspase 3 (CC3, brown) and periodic acid counterstain (pink). Glomerular apoptotic cells in aged IgG2a injected mice are indicated by the black arrows (B). (D) Quantification of the percent of glomeruli with a CC3 positive cell were increased in aged IgG2a injected mice (red bar) compared to young mice (gray bar) and were lowered with aPD1ab treatment (blue bar). (E-H) Representative images of immunoperoxidase staining for the ER stress marker GRP94 (brown). (E-G). (H) Quantification of GRP94 shows higher staining in aged IgG2a injected mice (red bar) which was lowered in IgG2a mice (blue bar). Individual mice represented by circles. (I-M) Autophagic activity measured by LC3. (I-K) Representative images of immunoperoxidase staining for microtubule associated protein 1 light-chain 3 (LC3) (brown). (L) Compared to young mice (gray bar), LC3 staining was lower in IgG2a injected mice (red bar) indicating reduced autophagy, but was higher with aPD1ab treatment (blue bar). (M) RNAseq data from isolated podocytes showed a decrease in the *Map1lc3a* (LC3) transcript in IgG2a injected mice (red bar) compared to young mice (gray bar) but was higher with aPD1ab treatment (blue bar).

This was confirmed by mRNA-seq data where transcripts for apoptosis genes (e.g., *Tp53*: 2-fold, *Tnf*: 1.35-fold, *Bim1*: 1.9-fold, *Card10*: 2.5-fold, and *Card14*: 1.8-fold) were elevated in aged control mice and reduced in the aPD1ab injected group. Interestingly, we also observed changes in pyroptosis, which is another form of podocyte death.(43) In podocytes from aged mice injected with the aPD1ab, inflammasome genes involved in pyroptosis were significantly downregulated compared to IgG2a control injected aged mice (e.g., *Nlrp2*: -2.72-fold, *Nlrp6*: -2.6-fold), *Nlrp4*: -2.56-fold, *Nlrp5*: -2.35-fold, *Gsdmd*: -2.11-fold, *Casp1*: -2.08-fold, and *Nlrp1b*: -2.59-fold).

Besides cell death, we looked at other forms of cellular stress such as endoplasmic reticulum stress (ERS) and autophagy.(44, 45) The staining intensity of the ERS associated proteins GRP94/Hsp90b1 increased in control IgG2a aged podocytes and was reduced in aged mice injected with the aPD1ab (**Figure 7E-H**). Conversely, the autophagy protein ATG8/MAP1LC3 (microtubule associated protein 1 light-chain 3 or LC3), a marker of autophagic activity, shows lower staining in glomeruli of control IgG2a aged mice (p<0.0001 *vs.* young) and is completely restored in aPD1ab-injected aged mice (p=0.001 vs aged control) (**Figure 7I-M**). Together, these data suggest that anti-PD-1 treatment of aged podocytes results in improved survival, reduced ER stress and augmented autophagy.

### Age-regulated Podocyte Signaling can be Restored by anti-PD-1 treatment

We have recently reported that the podocyte aging phenotype is caused by remarkable changes in autocrine and paracrine signaling molecules.(25) Thus, we wondered if anti-PD-1 injection altered this signaling network. Aged podocytes exhibit a striking inflammatory transcriptomic signature.(25) Hallmark pathway analysis (using a FDR cutoff of 0.05) showed that compared to age-matched IgG2a injected controls, aged mice injected with the aPD1ab exhibited a marked decrease in multiple inflammatory pathways (e.g. NOD-like, Toll-like receptor (TLR), interferon alpha and gamma, inflammatory genes, complement, allograft rejection, IL6-JAK-STAT, IL2-STAT, and KRAS) (**Supplementary Figures 6 & 7**). In addition, nine of the 52 ligand-receptor pairs regulated in aged podocytes(25) were impacted by aPD1ab injection (**Supplementary Figure 8**).

### Anti-PD1 antibody improves the podocyte’s metabolic state and reduced intracellular inflammation

Next, we performed a Gene Set Enrichment (GSEA) analysis of the mRNA-seq data to obtain a deeper understanding of the molecular pathways altered upon aPD1ab treatment. As shown in **Figure 8A&B**, the podocytes from mice injected with aPD1ab displayed decreases in T cell activation, positive regulation of protein kinase activity, mitotic nuclear division, DNA repair and calcium ion transport and marked increases in many metabolic pathways such as oxidative phosphorylation, amino acid metabolism, organic acid transport, mitochondrial translation and glucose metabolism. The latter was confirmed by Hallmark pathway analysis, which in addition to many metabolic pathways (e.g., oxidative phosphorylation, fatty acid metabolism, glycolysis, peroxisome) also identified regulation by the E2F transcription factors as GO terms upregulated in aged mice injected with aPD1ab compared to IgG2a (**Supplementary Tables**). Most impressively, overlaying those data onto individual pathways showed that aPD1ab injections caused up-regulation of many key components of Oxidative Phosphorylation, Glycolysis/Gluconeogenesis and the TCA cycle (**Figure 8C** and **Supplementary Figures 9 and 10**). Together these suggests that one of the primary activities of interfering with PD-1 signaling is a restoration of the metabolic profile of podocytes.

**Figure 8.**
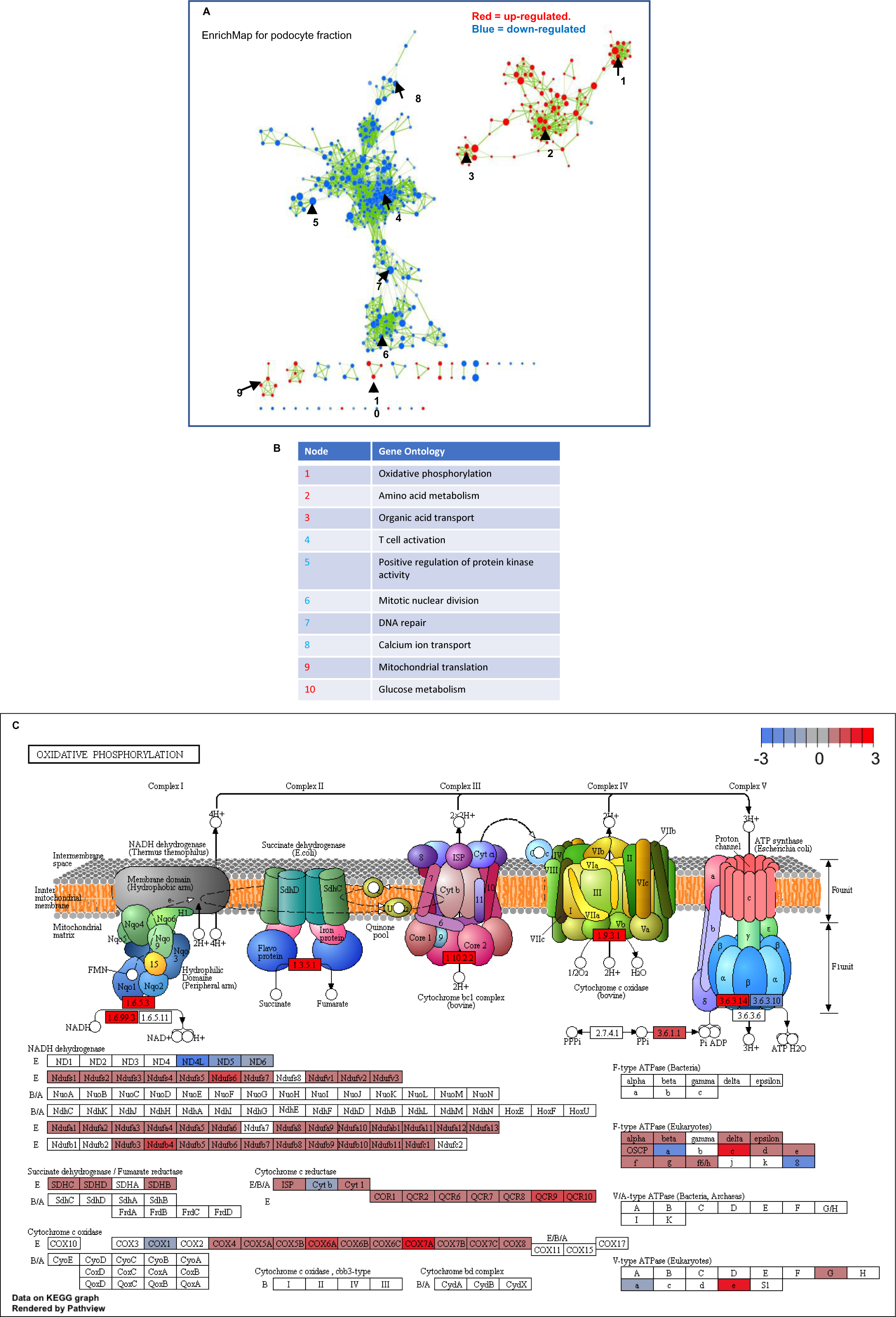
*Perturbed biological processes in podocyte aging and oxidative phosphorylation*. (**A**) Each node represents a gene ontology term significantly enriched in genes up-regulated in aged podocytes (red nodes) or down-regulated in aged podocytes (blue nodes). The transparency of the node is proportional to enrichment p-value: more solid-colored nodes represent more significantly enriched GO terms. Node size represent gene set size: GO terms with more genes have larger size. Two nodes are connected by an edge if they have overlapping genes. Edge transparency is proportional to the number of genes shared by two GO terms. Select GO terms from each cluster are labeled for each module. Overall, immune processes are up-regulated, while developmental and differentiation processes are down-regulated. Select nodes are numbered and their Gene Ontology terms are listed in the bottom right. (**B**) Table showing the names of the nodes for the pathways identified in A (**C**) Gene set enrichment for Oxidative Phosphorylation mapped to the KEGG pathway database. Schema shows genes that are decreased (blue) and increased (red) in the oxidative phosphorylation pathway in aged mice injected with anti-PD1 antibody (aPD1ab) compared to control aged mice given IgG2a. Scale in upper right shows fold change by color.

**Figure 9.**
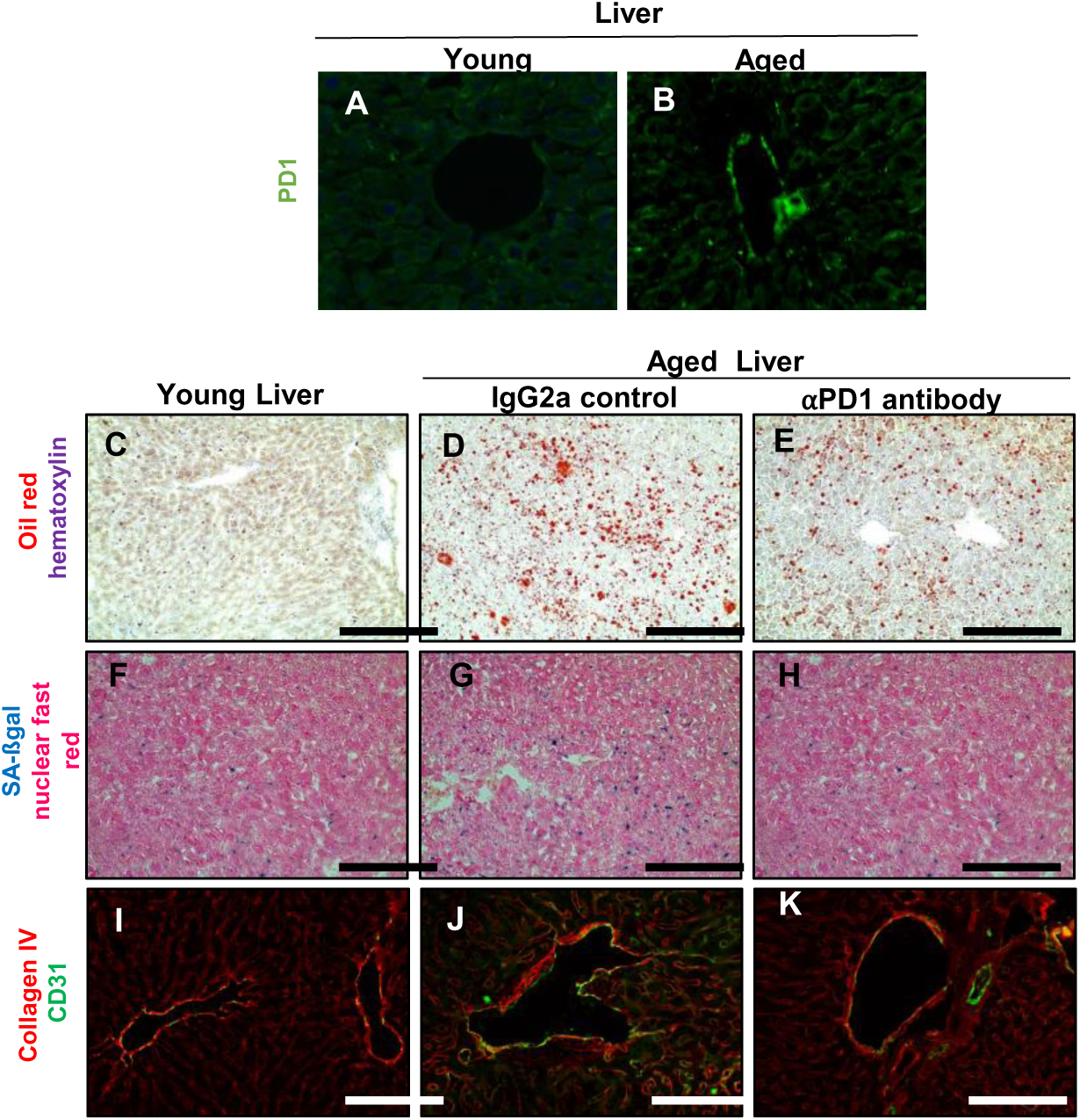
*Anti-PD1 antibody decreases liver aging in mice*. (A-B) Representative images of PD1 immunofluorescent staining which was higher in aged mouse liver (B) compared to young liver (A). (C-E) Oil red staining (red) used as a marker of fat deposition, was barely detected in young livers, but was increased in the livers of aged IgG2a injected mice and decreased in aPD1ab injected mice. (F-H) SA-ßgal staining (blue) used as a marker of senescence was increased in aged IgG2a injected livers and decreased by aPD1ab. (I-K) Double immunostaining for collagen IV (red) and the endothelial cell marker CD31 (green) shows increased collagen IV deposition in liver blood vessels in aged IgG2a injected mice, that was decreased by aPD1ab.

### Extra-glomerular effects of anti-PD-1 signaling on aging

Interfering with PD-1 signaling in mice is a systemic treatment and thus effects on cellular aging should not be restricted to the glomerulus. Indeed, tubular epithelial cell senescence assessed by SA-ßgal staining and immunostaining for the senescent proteins p16 and p19 was lowered upon aPD1ab injections (**Figure 3D-J**). In line with these observations, the mRNA-seq analysis of the non-podocyte fraction (i.e., the dissociated kidney cells remaining after MACs isolation of podocytes) showed upregulation of Gene Ontology terms such as amino acid metabolism, lipid catabolic process, oxidative phosphorylation and mitochondrial translation in the aPD1ab-injected kidneys as well as downregulation of processes such as epithelium morphogenesis, regulation of cell development, calcium ion transport, RAS protein signal transduction and leukocyte differentiation (**Data not shown**). Similarly, Hallmark pathway analysis shows decreases in EMT, TGFß signaling, several inflammatory pathways (e.g., TNA-alpha, NFKß, IL2-STAT5, complement signaling, IL6-JAK-STAT3), and hypoxia. As in the podocytes (**Figure 8**), many metabolic pathways were higher in aged mice injected with aPD1ab (data not shown). Immunostaining for Collagen IV and Interleukin 17A (Il17a) demonstrated that these changes were present both in the interstitium and the kidney epithelial cells, respectively (**Supplemental Figure 11**).

Finally, by extending the analysis to the liver, a similar anti-aging effect was observed outside of the kidney. PD-1 protein was present in the liver of aged mice, but not in young mice (**Figure 9A&B**).

Moreover, aPD1ab injection reverted the age-associated increase in liver fat deposition (oil red staining), senescence (SA-ßgal staining) and extracellular matrix deposition (Collagen IV immunostaining) when compared to control IgG2a-injected mice (**Figure 9C-K**). Together, these data suggest the PD-1 signaling is not restricted to the kidney and that aPD1ab injections have a more widespread benefit on aging phenotypes.

## DISCUSSION

Many tumors can avoid being detected by suppressing T cell immune responses upon activating negative regulatory pathways called immune check points. The latter include programmed cell death protein-1 (PD-1) and its ligands PD-L1 and PD-L2. The discovery of check point inhibitors for the PD-1 pathway prevents cancer cells from evading immune cells, thereby enabling successful T cell surveillance and subsequent killing of cancer cells.(29, 30, 46) However, much less is known about the PD-1 signaling pathway in non-tumor cells. Building on our recent report that PD-1 and its ligands are increased in aged mouse kidneys,(25) we now provide human data that glomerular *PCDC1* transcripts increase progressively with human aging, and the increase is statistically correlated with lower eGFR, higher segmental glomerulosclerosis and vascular damage. The current study asks if the glomerular consequences of increased PD-1 can be slowed or even reversed in the aged mouse kidney. We show that blocking PD-1 with an antibody improves the aging phenotype in kidneys and liver in mice with a particular improvement in both the life-span and health-span of aged podocytes. This interpretation is based on four major findings: (i) Aged mouse and human kidney display higher levels of PD1 immunostaining in epithelial cells (podocytes, PECs, tubular cells), but not in glomerular mesangial and endothelial cells. (ii) Ectopic expression of PD-1 in cultured podocytes triggers an apoptotic response. (iii) Interfering with PD-1 signaling in mice using a neutralizing anti-PD1 antibody reduces senescence markers in the kidney glomerulus, tubular epithelial cells and the tubular interstitium as well as the liver. (iv) While interfering with PD-1 signaling reverses some aspects of aging, it does not reverse all effects.

One interesting aspect of the study is interpreting the data in respect to health-span and life-span of podocytes, which are critical to the number and function of aging podocytes.(24) Injecting the aPD1ab to aged mice improves the life-span (i.e. survival) of podocytes by decreasing several pro-apoptotic and pyroptosis genes and increasing survival genes. At the same time, PD-1 inhibition also restores many of the features downregulated in aging podocytes required for their normal performance (health-span). These include not only individual canonical genes of functional podocytes (e.g., *Nphs1*, *Nphs2*, *Synpo* & *Vegfa*) or signaling networks (e.g., FGF signaling), but entire podocyte gene regulatory networks as illustrated by the VIPER analysis. Likewise, PD-1 inhibition improves the detrimental inflammatory phenotype present in aged podocytes which will further prevent podocyte dysfunction. Restoring these will obviously significantly impact the health-span of the podocyte and probably has a more long-lasting effect than simply preventing the death of individual podocytes. In this line, the observation that aPD1ab injections restores normal metabolic pathways such as oxidative phosphorylation, glycolysis and lipid metabolism in aged podocytes further strengthens the pro-health-span effects caused by PD-1 signaling inhibition. There may also be other consequences on improving the health of podocytes. For example, we have recently reported that aging podocytes not only exhibit autocrine loops, but also paracrine loops, in which podocytes signal to the other cell types in the glomerulus.(25) In the current study we observed that interfering with PD-1 signaling did not impact the age-dependent changes in glomerular mesangial or endothelial cell numbers, but injection of aPD1ab reduced the activated PEC phenotype (decreased CD44, CD74 and pERK) and the stressed GEN phenotype (decreased PV1). It is, thus, tempting to speculate that this “restoration” occurs via paracrine loops emanating from podocytes restored by the aPD1ab treatment. While our data on VEGFa as a surrogate marker for podocyte-endothelial crosstalk also support this hypothesis, more wide-ranging experimental analysis will be needed to address this in the future.

Studies have shown an association with increased PD-1 pathway expression and aging. The PD-1/PD-L1 pathway has been shown to be increased in aged dendritic aged cell subtypes and T cells,(47, 48) CD4+ T cells with features of cellular senescence,(49) and increases further with age in many cancers.(50)The PD-1/PD-L1 pathway is also increased in several kidney diseases independent of age, but are typically restricted to immune cells (reviewed in(51)) Kidney interstitial dendritic cells and human primary kidney proximal tubular epithelial cells express PD-L1 and PD-L2, where PD-L1 is integral for CD8 T cell tolerance.(52) Noteworthy is that several glomerulopathies have been recorded as a complication of anti-PD-1 immunotherapy.(52) We show for the first time that increased expression of *PDCD1* in aging human glomeruli is highly clinically significant, correlating with lower kidney function, higher scarring and increased vascular damage.

The study and usage of senolytics i.e., drugs that selectively clear senescent cells from an aging patient/organ is a vibrant field.(53) For example, the cancer drugs Quercetin and Dasatinib, which are broad spectrum inhibitors of protein kinases and tyrosine kinases have shown to reduce markers of aging.(54–56) Indeed, Dasatinib and Quercetin improve kidney function, increased expression of WT1 and decreased p16 levels in a diabetic model of senescence.(57) Similarly, induction of SASPs in aging cells has been a much discussed anti-aging target.(58) Yet, the heterogeneity of the SASP cytokines and its differences from cell type to cell type has made it difficult to develop viable anti-aging therapies. Thus, the focus has been towards some shared intracellular targets. Yet, our study now suggests that there may be an alternative avenue. Interfering with PD-1 in aged kidneys reduces the SASP of aging podocytes by markedly decreasing inflammatory pathways. Thus, upstream regulation of SASP may be a future avenue for anti-aging therapies. Obviously, caution needs to be considered because checkpoint inhibitors have been reported to have adverse effects in cancer immunotherapy causing renal toxicity.(59–61) Moreover, as in our study, systemic administration of the anti-PD1 antibody likely has off-target effects, and is not cell type-specific. While our data demonstrate good efficacy in the liver, other organs may respond differently. As such, integration of e.g., PD-1 inhibitors into anti-aging approaches will likely require a better understanding of the interaction between PD1 signaling and SASP induction.

We recognize the limitations of the current study, which include that anti-PD1 antibody treatment was given systemically rather than in a cell-specific targeted manner and was only given for a short duration. Yet, the impact on the kidney was impressive. Only one mouse species was studied, and undertaken in males, and the impact on organs beyond the kidney and liver were not assessed in this study. Finally, we did not explain the increase in PD-1 in aged podocytes. We speculate that the PD-1 transcriptional activator NF-kB is a major driver, as this is markedly increased in aged podocytes.

In summary, we show that the PD-1 pathway is increased in aged mouse podocytes and in human glomeruli. An increased expression of this pathway in human glomeruli predicts poor human kidney outcomes. Increased PD-1 levels in podocytes causes apoptotic loss (shortened podocyte life-span) and reduces their health-span. These events and glomerulosclerosis are partially reversed by inhibiting PD-1. These results are consistent with PD-1 being a mechanism contributing to glomerular damage in the aged kidney.

## METHODS

### Administering an anti-PD1 antibody to aged mice

21 month-old male C57BL/6J mice (∼ 70 year-old humans)(62)(63) were randomized into two groups: (1) anti-PD-1 antibody-injected group (n=12) that received weekly IP injections of InVivoMAb anti-mouse PD-1 antibody (RMP1-14, Bio X Cell, Lebanon, NH) at 12 mg/kg for 8 weeks; (2) aged-matched control mice (n=7) that received weekly IP injections of the isotype control IgG2A antibody (2A3, Bio X Cell, Lebanon, NH) at 12 mg/kg for 8 weeks. A third group of young uninjected C57BL/6J mice aged 4 months (n=9) were used as a non-aged comparison. Animals were weighed weekly; spot urines were collected prior to injections, at 4, and 8 weeks; albumin was measured by radial immunodiffusion assay (RID) and urine creatinine was determined with a Creatinine (urinary) Colorimetric Assay Kit (Cayman Chemical, Ann Arbor, MI).(64) Blood Urea Nitrogen was measured at sacrifice, with a colorimetric Urea Assay Kit (Abcam, Cambridge, United Kingdom).

### Immunostaining, Quantification and Visualization

Immunoperoxidase staining was performed on 4μm thick formalin fixed paraffin-embedded (FFPE) mouse and human kidney sections as previously described.(65) Double immunostaining was performed on 4 μm thick frozen and FFPE sections as previously described.(66) These immunohistochemical studies were approved by the University of Chicago institutional review board. The primary antibodies used in the study are summarized in **Supplemental Table 1**. To quantify immunohistochemistry, slides were scanned in brightfield with a 20x objective using a NanoZoomer Digital Pathology System (Hamamatsu City, Japan). The digital images were imported into Visiopharm software (Hoersholm, Denmark) and its Image Analysis Deep Learning module was trained to detect glomeruli and assess immunohistochemical staining-positivity for e.g., Collagen IV, p57 or ERG. The glomeruli ROIs were processed in batch mode generating per area outputs, cell counts and analyzed from 100% of the tissue sections. In the case of the parietal epithelial cells, images were collected using an EVOS FL Cell Imaging System (Life Technologies). Bowman’s capsule length was measured by using ImageJ 1.46r software (National Institutes of Health) and the percentage of positivity were calculated by dividing it by Bowman’s capsule length.

Two–dimensional images were detected on EVOS FL Cell imaging system (Thermo Fisher Scientific, Waltham, MA, USA) using 200x magnification. Stained kidney sections were digitally imaged by the University of Washington Histology and Imaging Core (HIC) using a Hamamatsu whole slide scanner (Bridgewater, NJ, USA).

### Assessment of liver aging

Fresh liver biopsies were preserved by snap freezing in OCT and stored at -80 C. Frozen sections were cut (8 μm thick) and used for staining as prescribed above. Hepatic lipids and triglycerides were determined by Oil Red O staining as previously prescribed.(67) To determine morphological changes present in aged liver, frozen liver sections (6μm) were stained for Collagen IV and the endothelial marker CD31/PECAM-1 as previously prescribed.(39)

### Magnetic activated cell sorting (MACS) of podocyte and non-podocyte cell fractions

Kidney tissue (w/o the kidney capsule and surrounding fat) was placed into ice cold RPMI 1640 medium, (w/o L-glutamine and phenol red, GE Healthcare Bio-Sciences, Pittsburgh, PA). After removal of the medulla, the remaining cortex was minced into fine pieces and digested in 0.2 mg/ml Liberase™ TL (Sigma-Aldrich, St. Louis, MO), 100 U/ml DNAse I (Sigma-Aldrich, St. Louis, MO) in RPMI 1640 medium (w/o L-glutamine and phenol red) by shaking at 37°C for 30 minutes. The digest was passed through an 18G needle (Becton Dickenson, Franklin Lakes, NJ) 10 times and enzymes were inactivated by adding 5 ml of RPMI 1640 medium (w/o L-glutamine and phenol red) supplemented with 1 mM sodium pyruvate (ThermoFisher Scientific, Waltham, MA), 9% Nu-Serum™ IV Growth

Medium Supplement (Corning Incorporated - Life Sciences, Durham, NC) and 100U/ml Penicillin-Streptomycin (ThermoFisher Scientific, Waltham, MA). The cell suspension was passed through a 100 µm and a 40 µm cell strainer (BD Biosciences, San Jose, CA) and pelleted by centrifugation at 200G at 4°C for 5 minutes. Cells were were resuspended in media containing two rabbit anti-Nephrin antibodies (1) (68) (1:100, Abcam, Cambridge, MA). After 1 hour at 4°C cells were pelleted, washed in media and incubated with anti-rabbit microbeads (Miltenyi Biotec, Auburn, CA) along with Alexa Fluor^®^ 594-conjugated AffiniPure Donkey Anti-Rabbit IgG 1:200 (in order to visualize binding of the Nephrin antibodies to the podocytes) for 30 minutes at 4°C. Cells were pelleted and washed in PBS with 0.5% BSA and 2mM EDTA and applied to MACS LS columns (Miltenyi Biotec, Auburn, CA) to gently separate microbead-bound podocytes from the other kidney cells. Cells not retained by the magnetic field were collected, pelleted and designated non-podocyte (NP) fractions. LS columns were removed from the magnetic field then washed with PBS with 0.5% BSA and 2mM EDTA and podocytes were collected. A small aliquot of each fraction were imaged using an EVOS FL Cell Imaging System to verify podocyte isolation, based on the presence of Nephrin antibody. Additionally, qRT-PCR for a panel of podocyte genes was performed in both podocyte and non-podocyte fractions to confirm cell type identity.

### RNA isolation, qRT-PCR, library preparation and sequencing

mRNA was isolated using the RNeasy Mini Kit (Qiagen, Germantown, MD) as per the manufacturer’s instructions and used for bulk mRNA sequencing or converted to cDNA by reverse transcription with the High-Capacity RNA-to-cDNA Kit (Thermo Fisher, Waltham, MA) and utilized for quantitative real-time PCR analysis. qRT-PCR was performed using iTaq SYBR Green Supermix (Bio-Rad, Hercules, CA) and a QuantStudio 6 Flex real-time PCR System (Applied Biosystems) as we have previously described.(25) Relative mRNA expression levels were normalized to glyceraldehyde-3-phosphate dehydrogenase (*Gapdh*). Library generation and bulk next-generation mRNA sequencing was performed by Psomagen, Inc. (Rockville, MD, USA) using TruSeq RNA Library Prep Kits (Illumina, San Diego, CA, USA) and the Illumina platform. Data analysis of the mRNA-seq data was performed as follows:

(1) Raw reads from fastq files were aligned to mm10 using the SubRead aligner.(69) Gene-level read counts were obtained using htseq-count.(70) Genes with less than 10 normalized reads summed across all samples were removed from further analysis. DESeq(71) was used to identify differentially expressed genes (DEGs), which were defined as genes with false discovery rate <0.05 and >2-fold change comparing young to aged. We additionally required that a DEG should have mean expression above 4 RPKM in either young or aged mice. Raw and processed RNA-seq data has been deposited at the NCBI Gene Expression Omnibus (accession number GSE186534 and can be browsed at the following link: https://yuliangwang.shinyapps.io/podocyte_PD1_experiment/
(2) Gene Set Enrichment Analysis (GSEA) was used to identify perturbed biological processes.(72) Genes were first ordered based on the pi score, a metric that combined p-value and fold change.(73) Genes with both a large increase in expression in aged podocytes and high statistical significance were at the top of the list; genes with both a large decrease in expression and high statistical significance were at the bottom of the list. Genes with moderate expression fold change and/or statistical significance were ranked in the middle. This ranked gene list was then used as input for GSEA.
(3) Additionally, the top Gene Ontology (GO) package(74) was used for GO enrichment analysis, based on Fisher’s exact test. In order to visualize the interactions between genes within perturbed pathways, we mapped DEGs onto network diagrams from the KEGG pathway database(75) using the PathView tool.(76)
(4) The VIPER (<u>v</u>irtual inference of <u>p</u>rotein activity by <u>e</u>nriched regulon analysis) method was used to identify potential master transcriptional regulators of podocyte aging as we described previously.(41) Genes are sorted from the most down-regulated genes in aged podocytes to the most up-regulated genes in aged podocytes compared to young podocytes. If a significant fraction of a TF’s positive targets were up-regulated, and its negative targets down-regulated in aged podocytes, this TF is inferred to be activated in aged podocytes (inactivated in young podocytes); if a significant fraction of a TF’s positive targets are down-regulated and its negative targets are up-regulated in aged podocytes, the TF is inferred to be inactivated in aged podocytes (activated in young podocytes).

### PD-1 overexpression in mouse podocytes

The Lenti-X 293T Cell Line (Takara Bio USA, Inc., Mountain View, CA) was cultured according to the manufacturer’s instructions and transfected with 5 μg pLenti-C-mGFP-P2A-Puro (control) or pLenti-C-mGFP-PD1-P2A-Puro (*Pdcd1* Overexpression), 5 μg of pLenti-Envelop vector and 6 μg of packaging plasmids (OriGene Technologies, Inc. Rockville, MD) to isolate lentivirus according to the manufacturer’s instructions. Immortalized mouse podocytes were isolated and cultured as previously described in detail (31–33) and infected with either pLenti-GFP control or pLenti-PD1 viral particles for 7 days and selected for expression using GFP expression via FACS. To confirm PD1 overexpression, mRNA was harvest and qPCR performed as described above. To block PD1, InVivoMAb anti-mouse PD-1 antibody (RMP1-14, Bio X Cell, Lebanon, NH) was applied every other day at 10ug/ml culture media for 5 days. To measure podocyte death, 40x or 100x random images were captured using an EVOS FL Cell Imaging System (Life Technologies) from 6 replicate samples per experimental group. Utilizing EVOS FL Auto 2 Software, the auto count feature was used to identify and count dead cells.

### Human data

Kidney tissue was obtained from the unaffected parts of kidneys removed from patients undergoing surgery at the University of Michigan and processed via the tissue procurement service of the Department of Pathology. Clinical data were obtained through the honest broker office of the University of Michigan as we have reported.(77, 78) Tissue was placed right away in RNAlater, micro-dissected into glomeruli and tubule-interstitial fractions, and isolated RNA was used for gene expression analysis using Affymetrix Human Gene 2.1 ST Array.(22, 78) This study was approved by the Institutional Review Board of the University of Michigan.

### Statistical analysis

Data are shown as the mean ± S.E.M. Student’s t-test was applied for comparisons between groups. Multiple groups were compared using one-way ANOVA with post hoc Tukey HSD test. P values <0.05 represented statistically significant differences.

### Study Approval

Animal studies were reviewed and approved by the University of Washington Institutional Animal Care and Use Committee (2968–04). Human studies were approved by the Institutional Review Board of the University of Michigan and University of Chicago institutional review board. Participants were provided written informed consent prior to participation.

### Author Contributions

Designed research studies: JWP, DGE, MB, OW & SJS: Conducted experiments: JWP, NK, YW, DGE, YZ, UT, CJL & COC; Acquired data: JWP, NK, YW, DGE, YZ, UT, CJL & COC, Analyzed data: JWP, NK, YW, DGE, COC, MB, OW & SJS; Wrote the manuscript JWP, NK, DGE, COC, MB, OW & SJS Provided material: AC

## Supporting information

DESeq

Hallmark Pathways

top Go Terms

DEGs up and down in aging via anti PD1

## Acknowledgements

Grants Supporting Work: 5 R01 DK 056799-10, 5 R01 DK 056799-12, 1 R01 DK097598-01A1, UC2 DK126006, R01 DK123031-01, NIH/NIA 5R01AG046231, DK12600601, W81XWH-19-1-0025

---

**Supplemental Figure 1.**
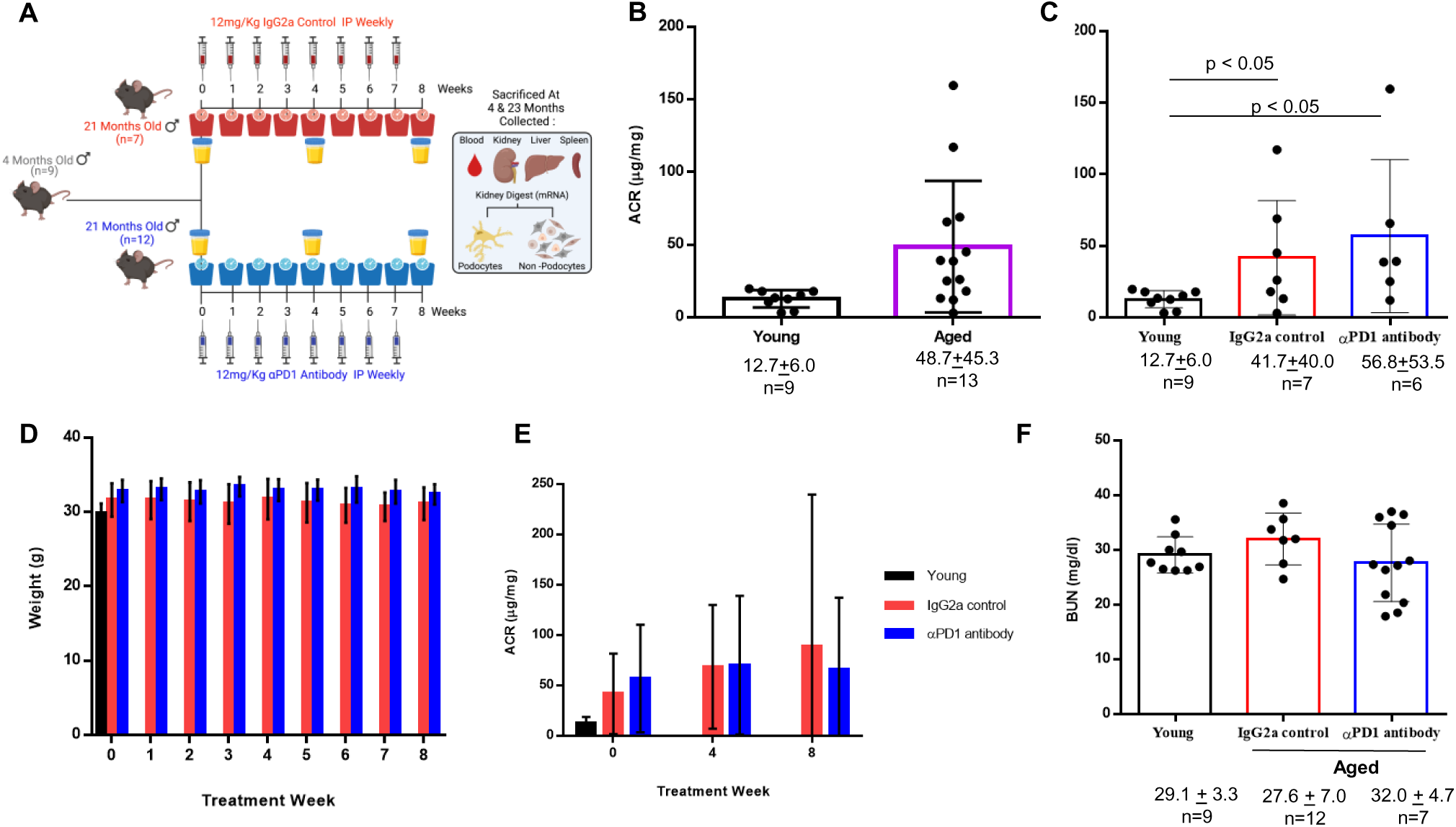
*Experimental Design and Physiological data for study animals*. (A) Schematic of study design. 4 month-old mice comprised the young group. The aged group of 21 month-old mice were randomized to the control group to receive 8 weekly intraperitoneal (IP) injections of IgG2a (control), or the treatment group to receive 8 weekly IP injections of anti-PD1 antibody. (B) Prior to randomization, uninjected aged mice (21m) have higher ACR versus uninjected young (4m) mice. (C) When randomizing the study groups, there was only a statistical difference (p<0.5) between young and aged animals, but not a difference between the two aged treatment groups. (D) For the duration of the study, aged animals did not lose weight with weekly injections of either the IgG2a (red) or anti-PD1 antibody (blue). (E) For the duration of the study, ACR values did not significantly change between or within study groups as a result of antibody injections. (F) BUN was not significantly different between young (4m) and aged IgG2a (23m), or anti-PD1 antibody (23m) injected mice.

**Supplemental Figure 2.**
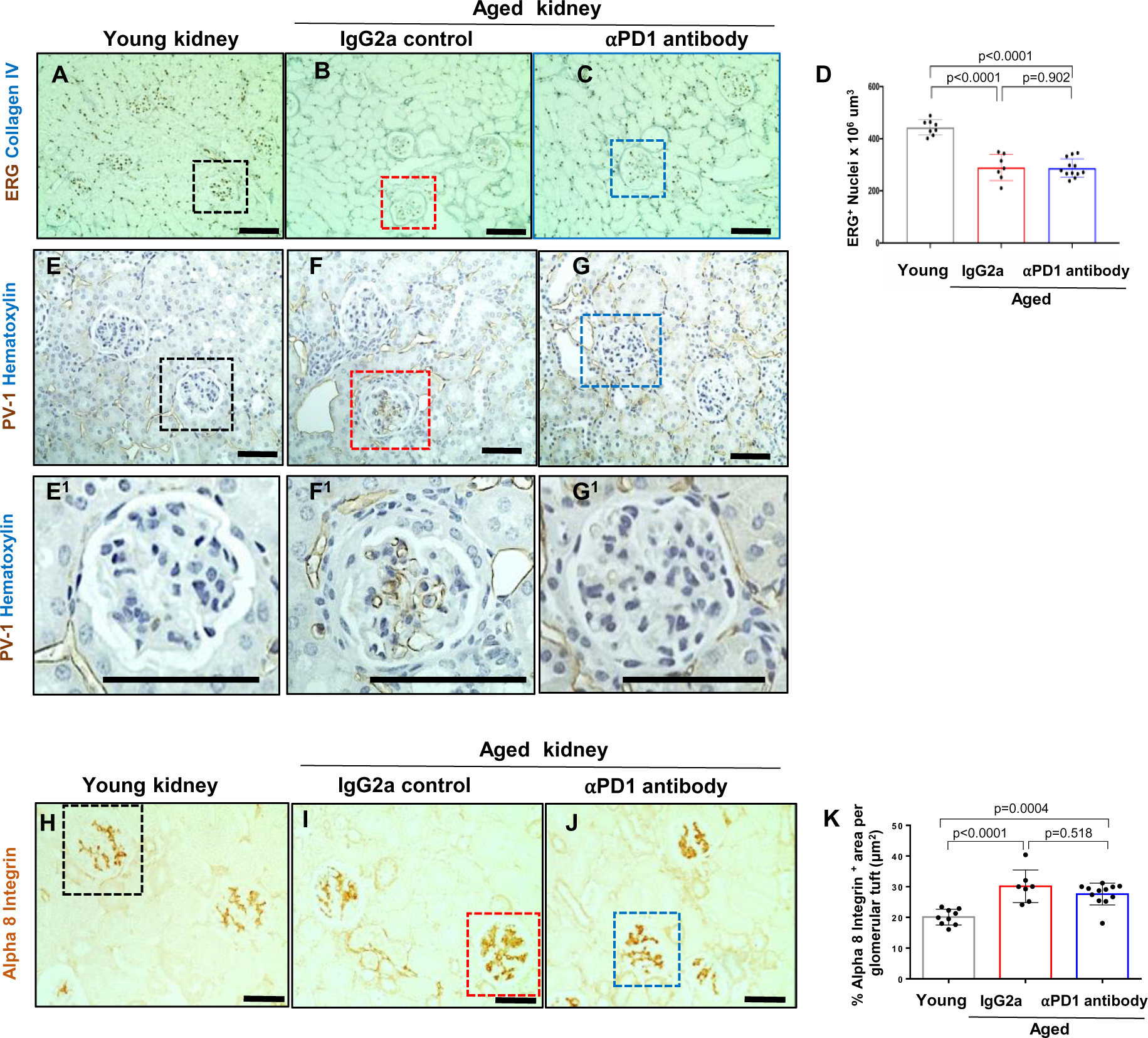
*Glomerular endothelial density/injury and mesangial cell density*. (A-C) Representative images of double staining for the glomerular endothelial cell marker ERG (brown, nuclear) and collagen IV (blue, outlines glomeruli). (D) Quantitation by AI shows a decrease in endothelial cell density in IgG2a injected mice, that does not change with aPD1ab treatment. (E-G) Representative images of immunoperoxidase staining for the fenestral diaphragm protein plasmalemmal vesicle associated protein-1 (PV-1) (brown) was not detected in glomerular endothelial cells of young mice, but was increased in aged IgG2a injected mice and decreased in aPD1ab injected mice. Superscripted images show higher magnification of the glomeruli highlighted by dashed boxes. (H-J) Representative images of immunoperoxidase staining for the mesangial cell marker alpha 8 integrin (brown). (H) Quantification of staining showed an increase in aged IgG2a injected mice that does not change with aPD1ab treatment.

**Supplemental Figure 3.**
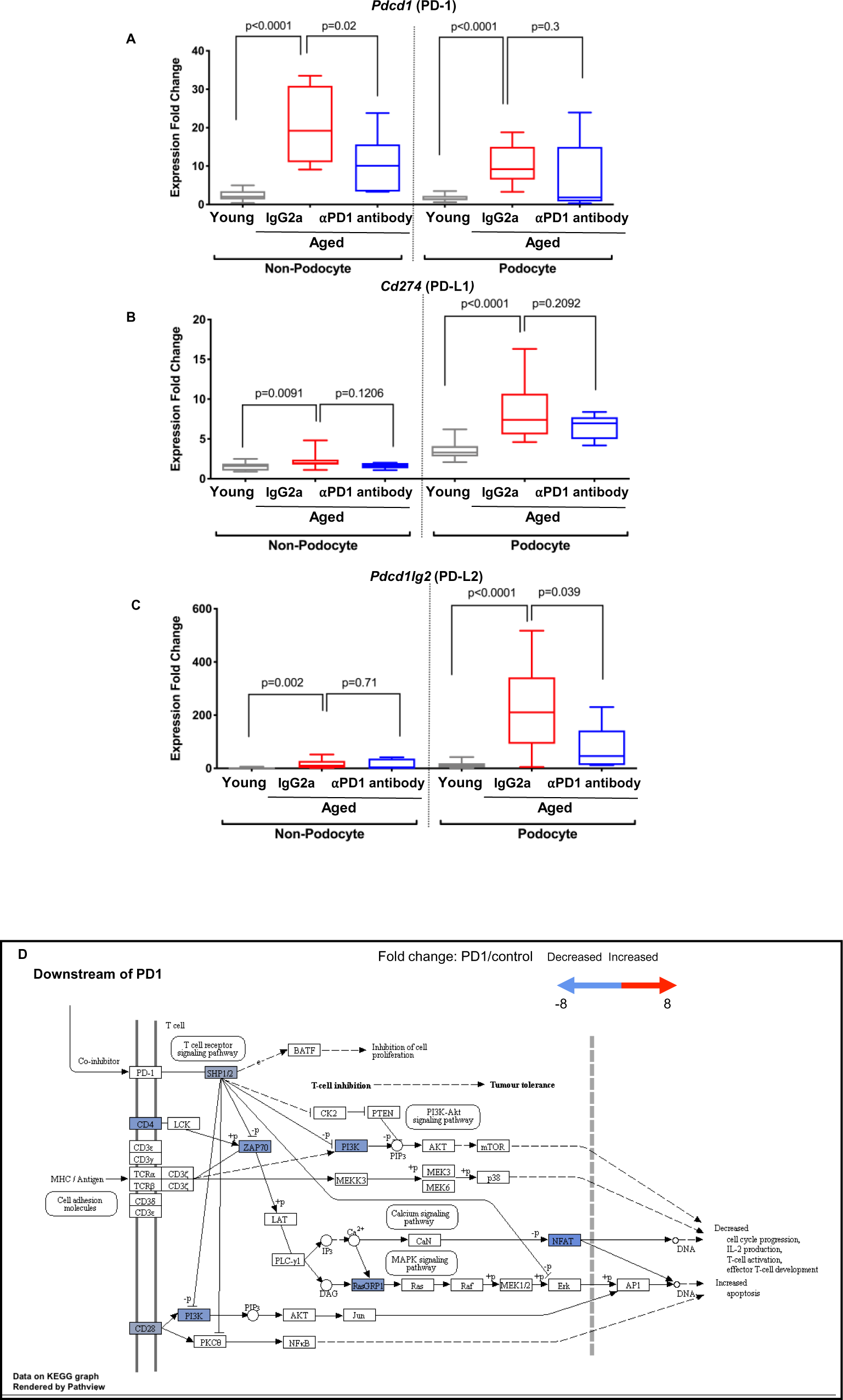
*Changes to the PD1 pathway in aged podocytes by anti-PD1 antibody*. (A-C) mRNA expression measured by qRT-PCR for *Pdcd1* (PD1), *Cd274* (PD-L1) and *Pdcd1lg2* (PD-L2) in non-podocytes and podocytes. In the non-podocyte fraction, aPD1ab did lower *Pdcd1*, but did not change levels of *Cd274* or *Pdcd1lg2* compared to IgG2a injected mice. PD-L1 and PD-L2. In podocytes, aPD1ab did appear to lower *Pdcd1*, *Cd274* and *Pdcd1lg2*, compared to IgG2a injected mice, but this did not reach statistical significance due to variability. (**D**) Gene expression data was mapped to PD1 target pathways in KEGG pathway database. Red rectangles denote up-regulated genes (aged/young expression ratio >1), blue rectangles denote down-regulated genes (aged/young expression ratio <1); grey rectangles denote genes with no significant change. Edges represent interactions between genes/proteins. “+p” denotes phosphorylation. Sharp arrows indicate positive, while dashed arrows indicate negative regulation. Scale in upper right shows fold change by color. The results showed an overall significant decrease in genes downstream of PD1 in aged podocytes from mice injected with aPD1ab compared to control aged-matched mice given IgG2a.

**Supplemental Figure 4.**
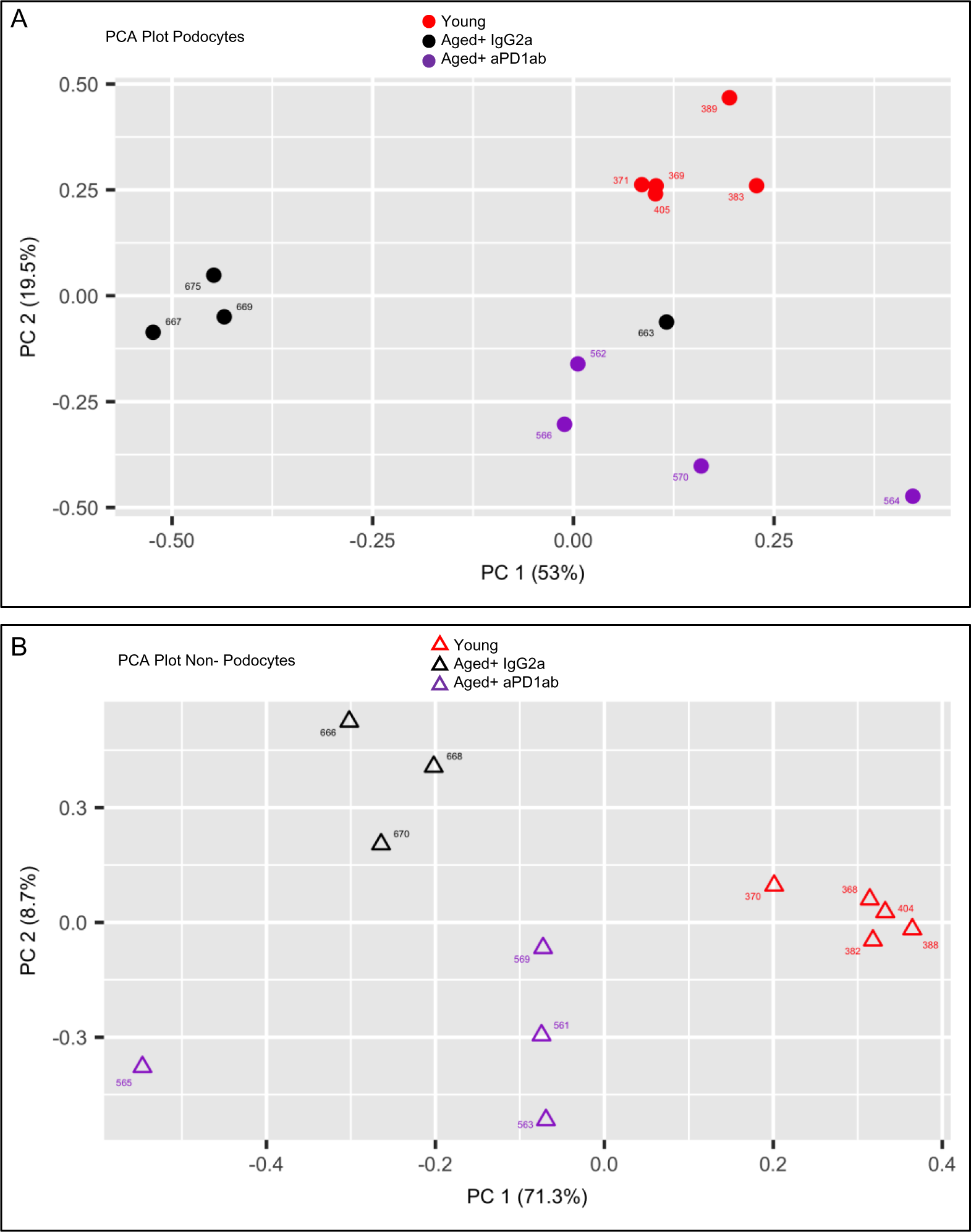
*Principal Component Analysis (PCA)*. Principle component analysis showed excellent clustering of the individual treatment groups in both podocytes (A) and non-podocytes (B)

**Supplemental Figure 5.**
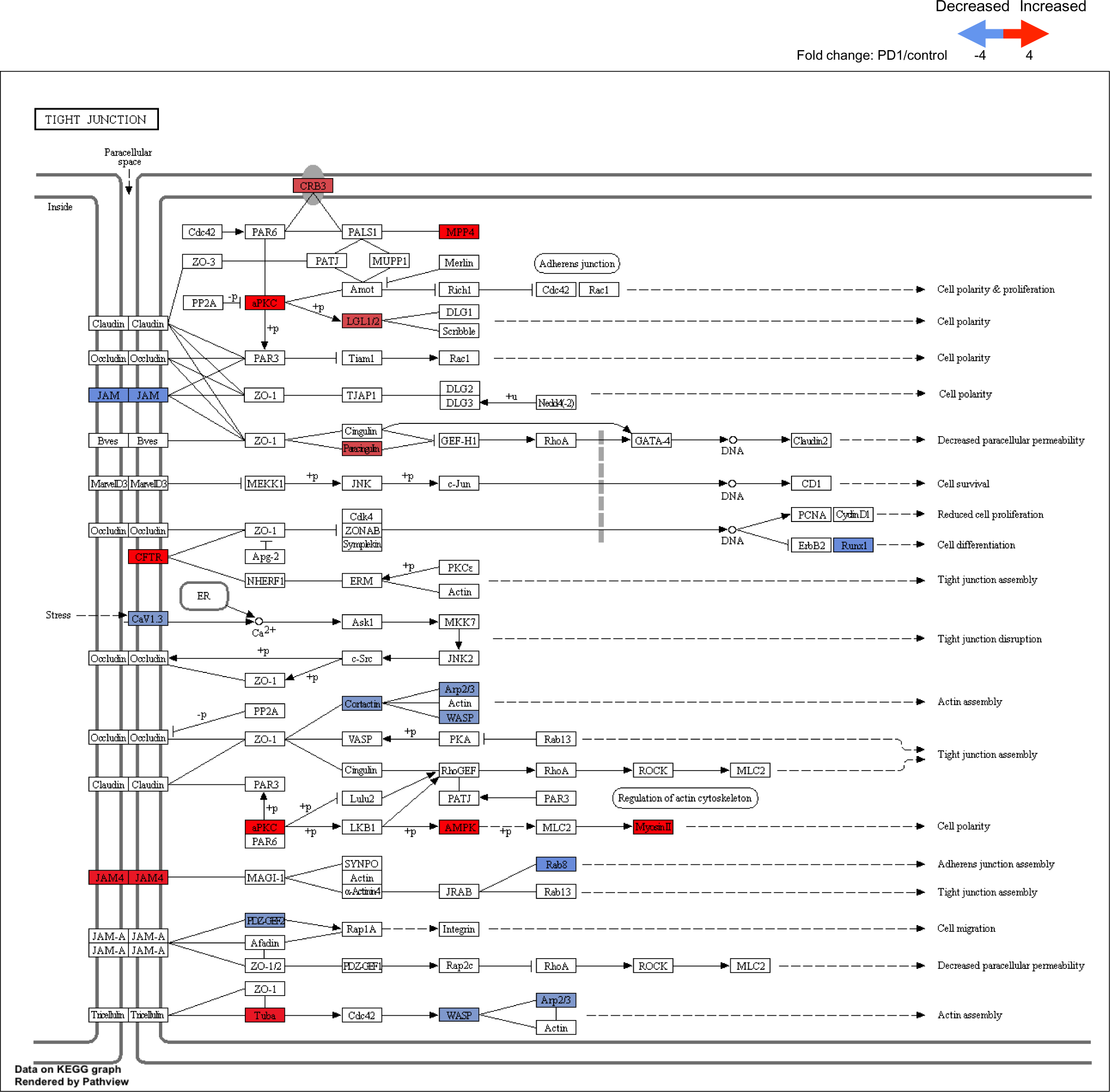
*Gene expression data mapped to the tight junction pathway in the KEGG pathway database*. Gene expression data was mapped to tight junction pathways in KEGG pathway database. Red rectangles denote up-regulated genes (aged/young expression ratio >1), blue rectangles denote down-regulated genes (aged/young expression ratio <1); grey rectangles denote genes with no significant change. Edges represent interactions between genes/proteins. “+p” denotes phosphorylation. Sharp arrows indicate positive, while dashed arrows indicate negative regulation. Scale in upper right shows fold change by color. The results showed an overall significant increase in genes for tight junctions in aged podocytes from mice injected with aPD1ab compared to control aged-matched mice given IgG2a.

**Supplemental Figure 6.**
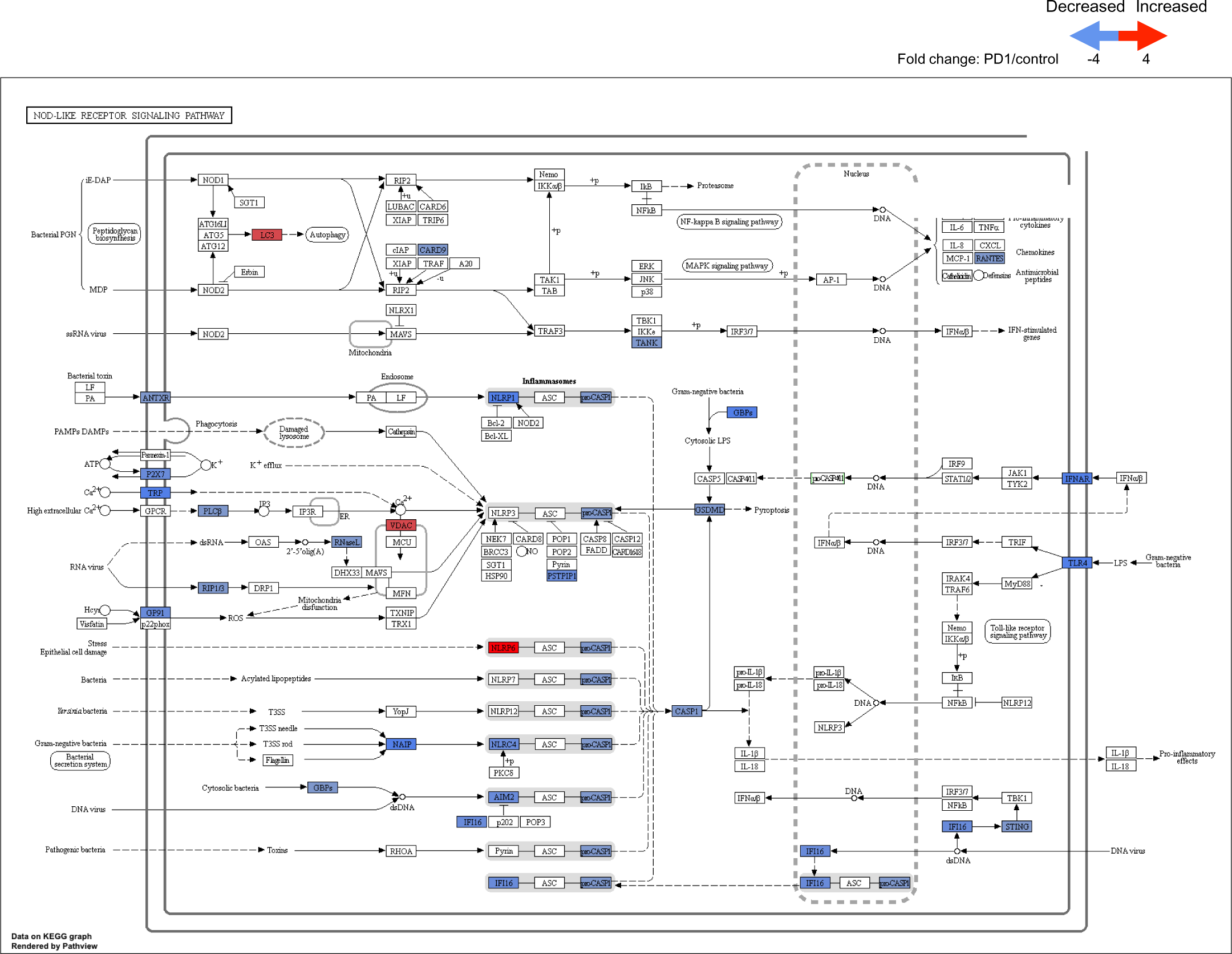
*Gene expression data mapped to NOD-Like Receptor Signaling Pathway in the KEGG pathway database*. Gene expression data was mapped to NOD-Like Receptor Signaling Pathway in KEGG pathway database. Red rectangles denote up-regulated genes (aged/young expression ratio >1), blue rectangles denote down-regulated genes (aged/young expression ratio <1); grey rectangles denote genes with no significant change. Edges represent interactions between genes/proteins. “+p” denotes phosphorylation. Sharp arrows indicate positive, while dashed arrows indicate negative regulation. Scale in upper right shows fold change by color. The results showed an overall significant decrease in genes for NOD-Like Receptor Signaling Pathway in aged podocytes from mice injected with aPD1ab compared to control aged-matched mice given IgG2a.

**Supplemental Figure 7.**
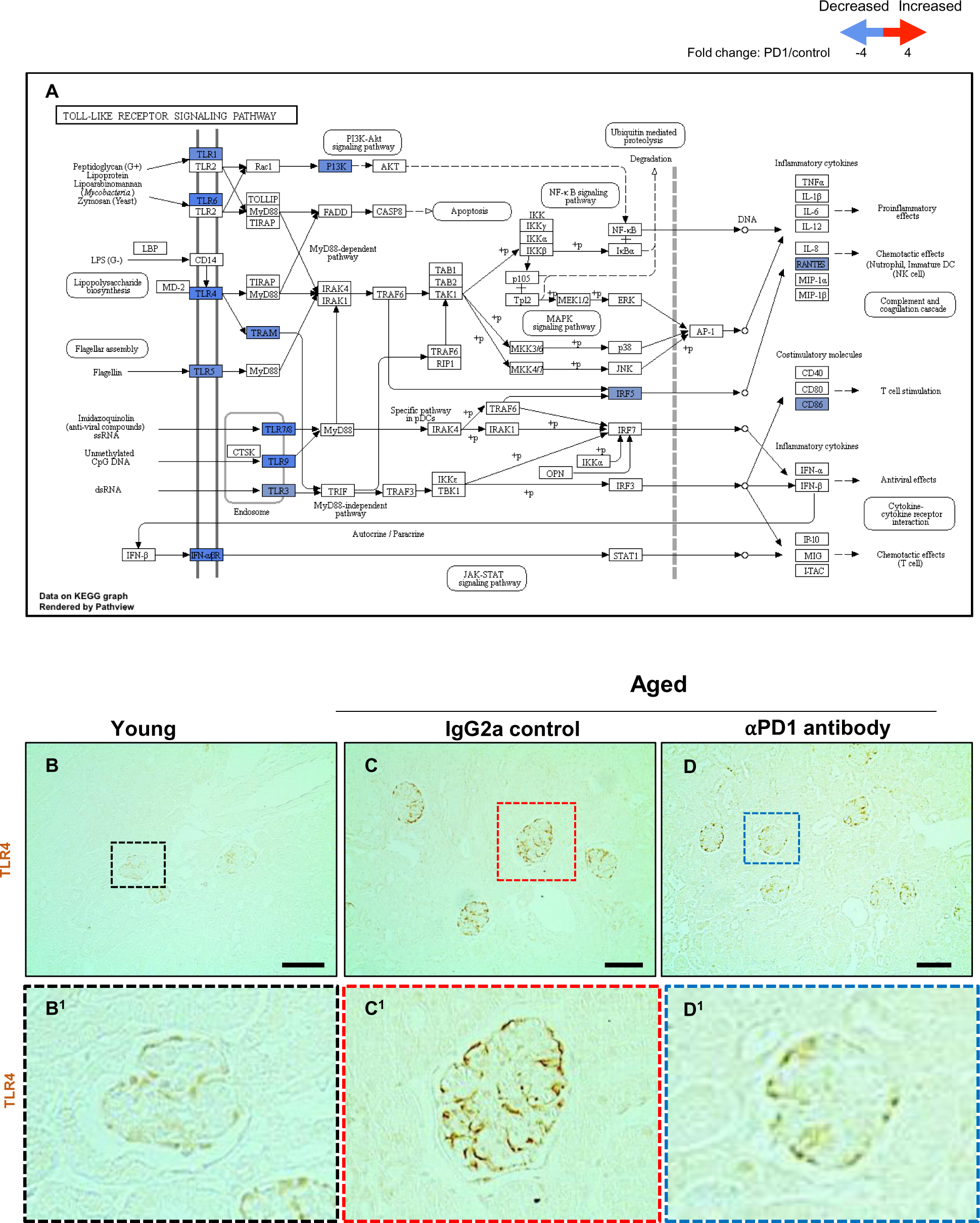
*Gene expression data mapped to Toll-Like Receptor Signaling Pathway in the KEGG pathway database*. (A) Gene expression data was mapped to Toll-Like Receptor Signaling Pathway in KEGG pathway database. Red rectangles denote up-regulated genes (aged/young expression ratio >1), blue rectangles denote down-regulated genes (aged/young expression ratio <1); grey rectangles denote genes with no significant change. Edges represent interactions between genes/proteins. “+p” denotes phosphorylation. Sharp arrows indicate positive, while dashed arrows indicate negative regulation. Scale in upper right shows fold change by color. The results showed an overall significant decrease in genes for Toll-Like Receptor Signaling Pathway in aged podocytes from mice injected with aPD1ab compared to control aged-matched mice given IgG2a. (B-D) Representative images of immunoperoxidase staining (brown) showing increased staining for TLR4 in a podocyte distribution in aged IgG2a mice (C) compared to young mice (B), which is lower in aged mice injected with aPD1ab (D). Panels with superscript show higher magnification of the glomeruli in the dashed insets.

**Supplemental Figure 8.**
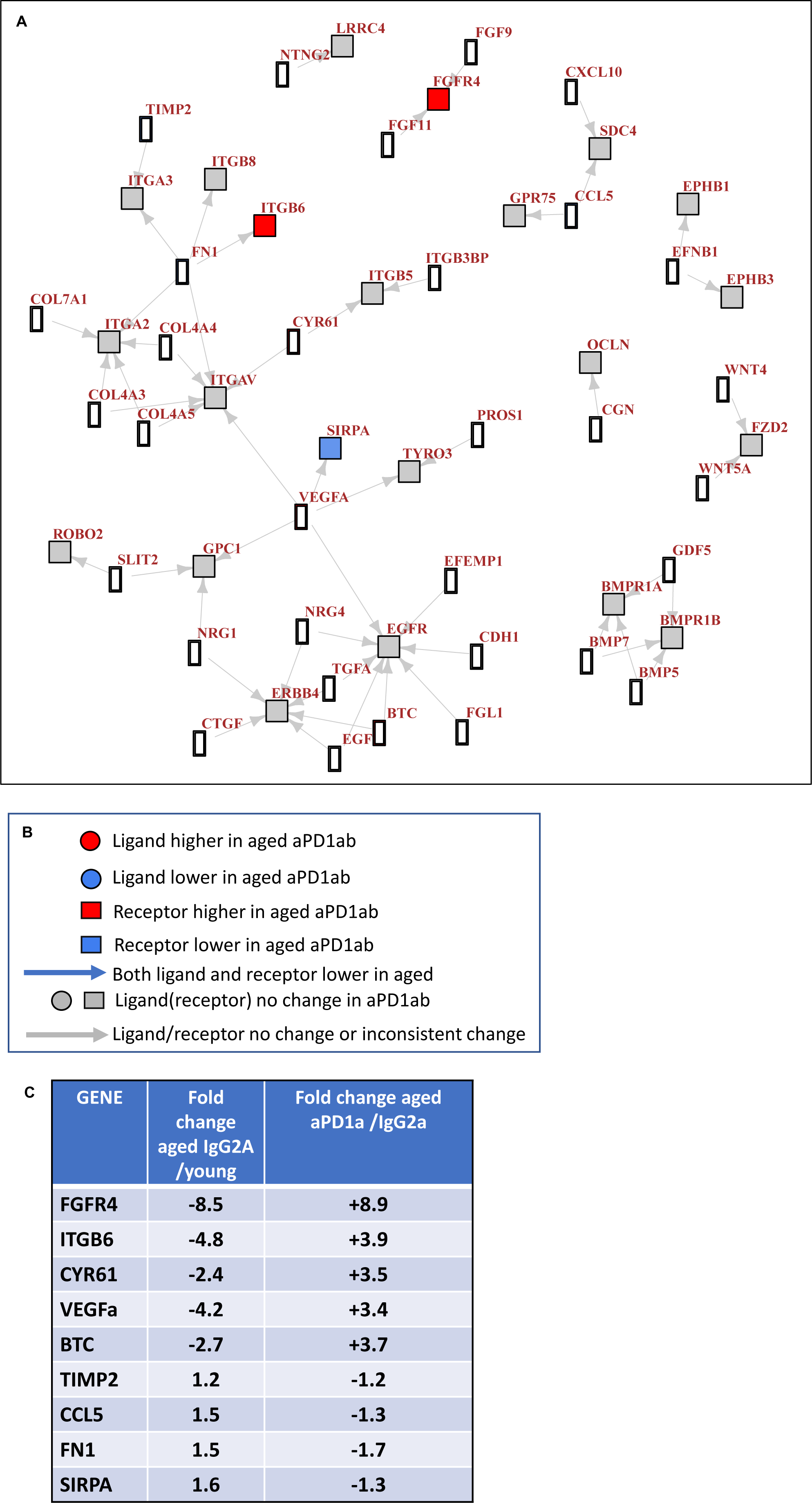
*Ligand-Receptor Analysis*. (A) Mapping of ligands with their receptors that were increased in the podocytes of aged IgG2a injected mice compared to young podocytes. (B) Legend showing changes in aged aPD1ab injected mice compared to aged IgG2a injected mice. (C) Fold changes for genes impacted by aPD1ab.

**Supplemental Figure 9.**
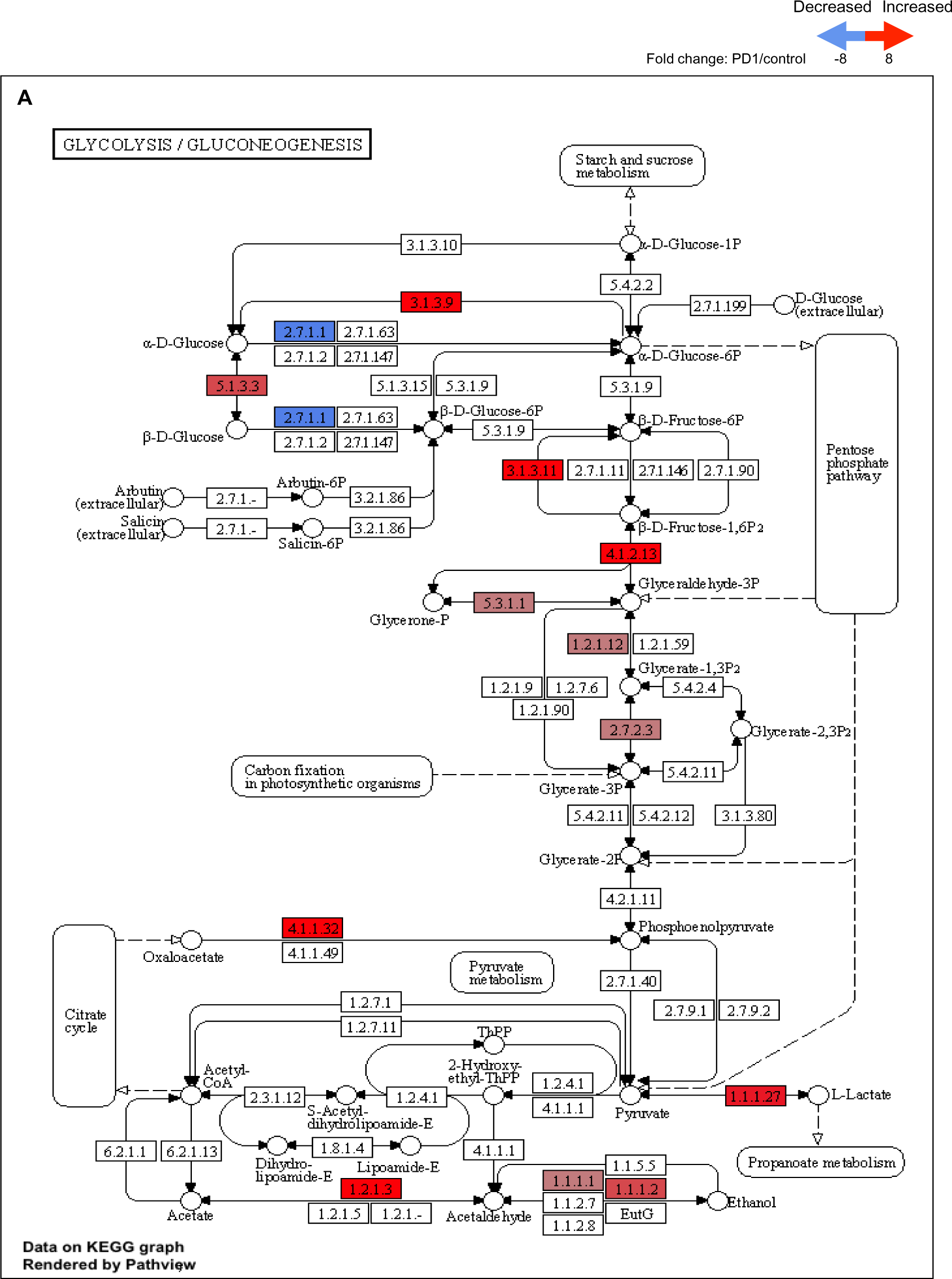
Gene expression data mapped to the Glycolysis/Gluconeogenesis pathway in the KEGG pathway database. (A) Gene expression data was mapped to the Glycolysis/Gluconeogenesis Pathway in KEGG pathway database. Red rectangles denote up-regulated genes (aged/young expression ratio >1), blue rectangles denote down-regulated genes (aged/young expression ratio <1); grey rectangles denote genes with no significant change. Edges represent interactions between genes/proteins. “+p” denotes phosphorylation. Sharp arrows indicate positive, while dashed arrows indicate negative regulation. Scale in upper right shows fold change by color. The results showed an overall significant increase in genes for to the Glycolysis/Gluconeogenesis Pathway in aged podocytes from mice injected with aPD1ab compared to control aged-matched mice given IgG2a.

**Supplemental Figure 10.**
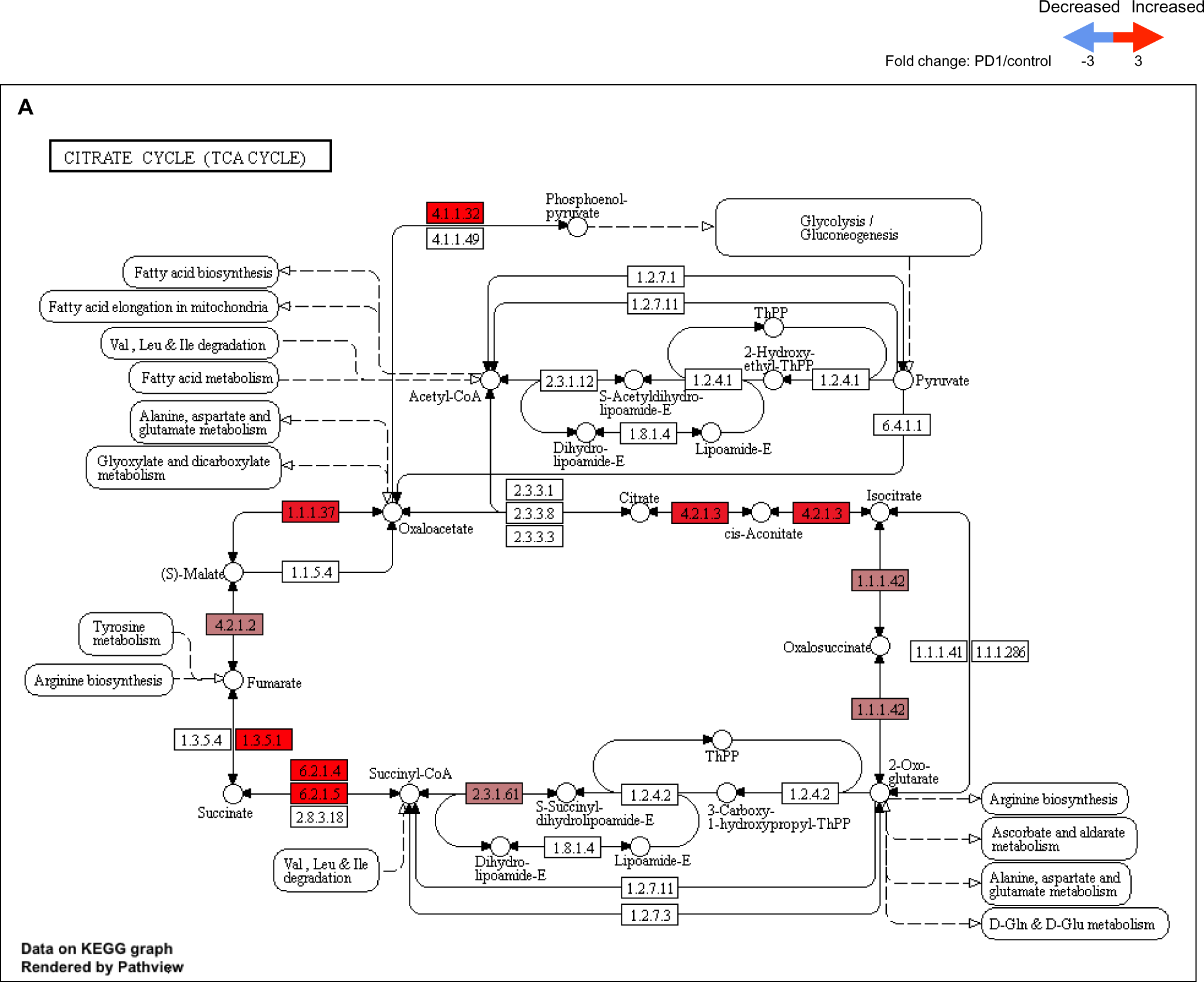
*Gene expression data mapped to the Citrate Cycle pathway in the KEGG pathway database*. (A) Gene expression data was mapped to the Citrate Cycle Pathway in KEGG pathway database. Red rectangles denote up-regulated genes (aged/young expression ratio >1), blue rectangles denote down-regulated genes (aged/young expression ratio <1); grey rectangles denote genes with no significant change. Edges represent interactions between genes/proteins. “+p” denotes phosphorylation. Sharp arrows indicate positive, while dashed arrows indicate negative regulation. Scale in upper right shows fold change by color. The results showed an overall significant increase in genes for to the Citrate Cycle Pathway in aged podocytes from mice injected with aPD1ab compared to control aged-matched mice given IgG2a.

**Supplemental Figure 11.**
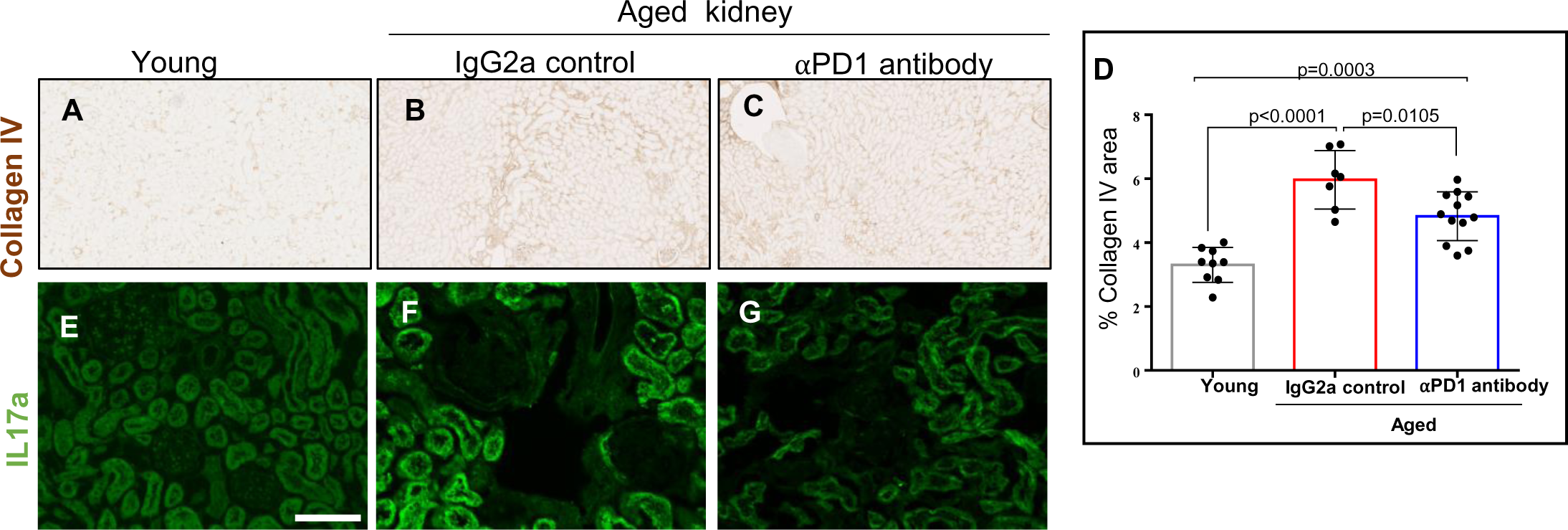
*Tubulointerstitial Changes with aPD1ab injection*. (A-C). Representative images of collagen IV immunoperoxidase staining (brown). Compared to young kidneys (A), Collagen IV staining was higher in the interstitium of aged IgG2a injected mice (B), which is decreased by aPD1ab (C). (D) Quantitation of collagen IV staining by image analysis. (E-G) Representative images of Interleukin 17a (IL17a) immunofluorescence (green) staining. Compared to young kidneys (E), IL17a staining was higher in the tubular epithelial cells in aged IgG2a injected mice and was decreased by aPD1ab (G).

